# TPR domain assigns versatility of BcTir/Tpr system against viral infection

**DOI:** 10.1101/2023.05.26.542521

**Authors:** Yingying Liu, Chen Zhang, Nelli Khudaverdyan, Junyi Wang, Lei Zhang, Mikhail Y Golovko, Svetlana A Golovko, Ang Guo, Jikui Song, Min Wu, Colin Combs

**Author notes:** Correspondence (M.W), (C.C). These authors contributed equally (Yingying Liu, Chen Zhang). retired (Min Wu).

## Abstract

NAD^+^-derived signal produced by TIR domain triggered the host immune responses. The ubiquitous TPR domain involved in signal recognition and effector activation were found widely assembled with the TIR domain. However, the immune roles of these assemblies remain elusive. Here, a two-gene operon, one containing a TIR domain, designated as BcTir, and the other, BcTpr, from *Bacillus cereus*, exhibited anti-phage immunity. BcTpr, but not BcTir, exhibited NADase activity to produce the cyclic ADPR (cADPR) isomer and mediate NAD^+^ depletion. Noticeably, the truncated N terminus of BcTpr only depleted NAD^+^ unless at the presence of TPR domain to generate cADPR isomer unveiling its role played for glycosite selection. In addition, the BcTir/Tpr system significantly repressed viral proliferation and increased oxidation resistance by scavenging excessive reactive oxygen species (ROS) upon phage infection. These findings unraveled a multifunctional role of the BcTir/Tpr system during immune responses.

**In Brief:** The bacterial BcTir/Tpr system was identified with the ability to protect against phage infection via NAD^+^ depletion, viral replication repression, and ROS homeostasis, in which BcTpr played a dual role in NAD^+^-derived signal production and NAD^+^ depletion.

**Highlights:**

- The BcTir/Tpr system works as a BcTir-BcTpr complex against phage infection.
- BcTpr, instead of BcTir, generates the NAD^+^-derived cADPR isomer through its glycosidase domain at N terminus.
- The amount of cADPR isomer production is regulated by the helix numbers of the TPR domain at C terminus of BcTpr.
- The BcTir/Tpr system can depress phage proliferation and decrease ROS production upon phage infection.

## Introduction

TIR-containing (Toll/interleukin-1 receptor domain) proteins or systems, such as SARM1 in humans, the nucleotide-binding NLR (leucine-rich repeat-containing) in plants, and the thoeris system, ThsA/ThsB, in bacteria, are characterized with NADase activity to generate NAD^+^-derived messengers, such as ADP-ribose (ADPR) or its cyclic variants that subsequently activate downstream effectors for depleting NAD^+^ to mediate immune responses, like cell death via abortive infection (Essuman et al., 2017; Essuman et al., 2018; Ka et al., 2020; Ofir et al., 2021; Tian et al., 2021).

In parallel, a ubiquitous domain, tetratricopeptide repeats (TPR), across the tree of life, is also a critical defense element responding to environmental changes, including viral infection and abiotic stresses via switching the bonded enzymatic activity on and off (Cui et al., 2022; van Beljouw et al., 2021; Weijman et al., 2017; Zhou et al., 2018). Moreover, the TPR domain is reported as essential for the recognition and selection of glycosides and guides the related enzymes to post-translationally modify thousands of protein substrates closely associated with many diseases such as neurodegeneration (Allan and Ratajczak, 2011; Ramirez et al., 2021). Recent genomic analyses identified a range of TPR-containing assemblies (Burroughs and Aravind, 2020), in which a few TPR domains were consistently neighbored by a TIR domain, suggesting coordination in guiding NAD^+^-related signal transduction and likely regulation of the antiviral responses. Therefore, their interconnection for immunity is worth elaboration.

Here, we set out to investigate the functional roles of a small operon representative from *Bacillus cereus* (OOQ93166.1) that consists of two genes, encoding BcTir and BcTpr, respectively, designated as the BcTir/Tpr system, in which TIR and TPR domains are separately located (Burroughs and Aravind, 2020). Our study identified that the BcTir/Tpr system armed the *E. coli* bacterium (BL21) with the ability to combat phage infection involving NAD^+^ metabolism. BcTpr, but not BcTir, utilized NAD^+^ and generated the cADPR isomer as a converted messenger. Moreover, the BcTir-BcTpr complex was a prerequisite for coordinating the bacterial defense against phage infection, which is distinct from the paradigm of TIR functioning to transform NAD(P)^+^ to (v)cADPR as previously reported (Essuman et al., 2022). In addition, the BcTir/Tpr system plays a multifunctional role in repressing viral replication and removing excessive ROS produced during phage infection, thus minimizing oxidation-associated damage by upregulating enzymatic scavengers. The BcTir/Tpr system exhibited pleiotropic functions to fortify bacterial host defense against invaders.

## Results

### Anti-viral immunity conferred by BcTir/Tpr system

We first sought to test the immunity potential of the BcTir/Tpr system from *Bacillus cereus*. Thus, the BcTir/Tpr system, was cloned and transformed into *E. coli* BL-21 (DE3) recipients (Figure 1A). Our initial experiments showed that expression levels of the BcTir/Tpr genes were elevated with increasing IPTG (Figure S1A). The bacterial growth was inhibited when IPTG reached concentrations of 50 μM or higher and to minimize the side effect of high concentration IPTG on cell growth (Dvorak et al., 2015; Einsfeldt et al., 2011), we elected to use 50 μM IPTG in experiments thereafter. T-series lytic phages, including T2, T3, T4, T5, and T7, and P1 as the temperate phages, were employed to investigate the bacterial immunity potency conferred by the BcTir/Tpr system against phage infection (Lopatina et al., 2020; Seed, 2015). Our results revealed that the BcTir/Tpr system strongly impeded viral proliferation of T phages, especially T2 phage, compared to control *E. coli* with an empty vector (Figure 1B). However, the expression of either BcTir or BcTpr alone was unable to prevent phage infection, supporting the notion of coordinated action between the two domains derived from the separate encoding genes are required for this ability. Similarly, the BcTir/Tpr system also prevented P1 phage infection but to a lesser extent. Thus, the T2 phage was selected for the downstream functional assays of the BcTir/Tpr system. We also noticed that expression of the BcTir/Tpr system was enhanced after T2 phage infection at 30 min at every IPTG concentration. This increase further supported a role for the BcTir/Tpr system against phage attack (Figure S1B). *E. coli* bearing the BcTir/Tpr system had no growth difference compared to transformants with an empty vector under non-phage infection (Figure 1C). T2 phage infection at a multiplicity of infection (MOI) of 2 showed that the BcTir/Tpr system significantly retarded cell growth arrest after 30 min of T2 phage infection versus the control *E. coli* with an empty vector (Figure 1C). Separately expressed BcTir or BcTpr followed a similar trend as the control and could not stop the cell death even at lowered MOIs (Figure S2). Furthermore, the cell growth arrest of *E. coli* containing BcTir/Tpr, not the separated BcTir or BcTpr, was markedly reversed when the MOI was decreased to 0.02, indicating that this system can prevent further cell destruction as confirmed by fluorescence activated cell sorting (FACS) (Figure S3). Similar to the ThsA/ThsB system (Ofir et al., 2021), the BcTir/Tpr system also depleted NAD^+^ completely after 15 min of phage infection, in which only the BcTpr but not BcTir (Figure 1D) aligned with the *in vitro* data (Figure 1E). Our analysis of the BcTir crystal structure at 1.78 Å resolution revealed a three-stranded parallel β-sheet, packed by two bridging α-helices on one side and another three on the other side, homologous to the TIR domain of human SARM1 (hSARM1) as well as the N-terminal subdomain of ThsB in the Thoeris system (Ka et al., 2020; Ofir et al., 2021) (Figure S4A). Coincidently, the active catalytic site, glutamate (E) of the TIR domain (hSARM1), was naturally mutated to lysine (K) at site 37 of BcTir (Figure S4B), and a site mutation, BcTir K37E, partially recovered the NAD^+^-depleting potential. In addition, BcTir K37E, BcTpr, and BcTir/Tpr were consistently expressed and maintained intracellular NAD^+^ at a lower level compared to the control without phage infection, which did not affect the correspondent growth of *E. coli*, consistent with recent observations (Coronas-Serna et al., 2020). Moreover, a product generated by BcTpr had the same molar mass as cyclic ADP-ribose (cADPR) detected by tandem mass spectrometry (MS/MS) (Figure S5), but with a different retention time, suggesting that this product is the NAD^+^-derived isomer of cADPR (Figure 1F). Subsequent measurements revealed that BcTpr, rather than BcTir, was responsible for producing the cADPR isomer, which was induced over three-fold in the BcTir/Tpr system at 15 min post phage infection compared to that of no phage infection (Figure 1G). By contrast, the level of cADPR isomer was maintained by BcTpr at high concentrations and at all time points before cell destruction after 45 min post phage infection, whereas it was repressed to low levels by the BcTir/Tpr system, indicating a regulatory role of BcTir for cADPR isomer production.

**Figure 1.**
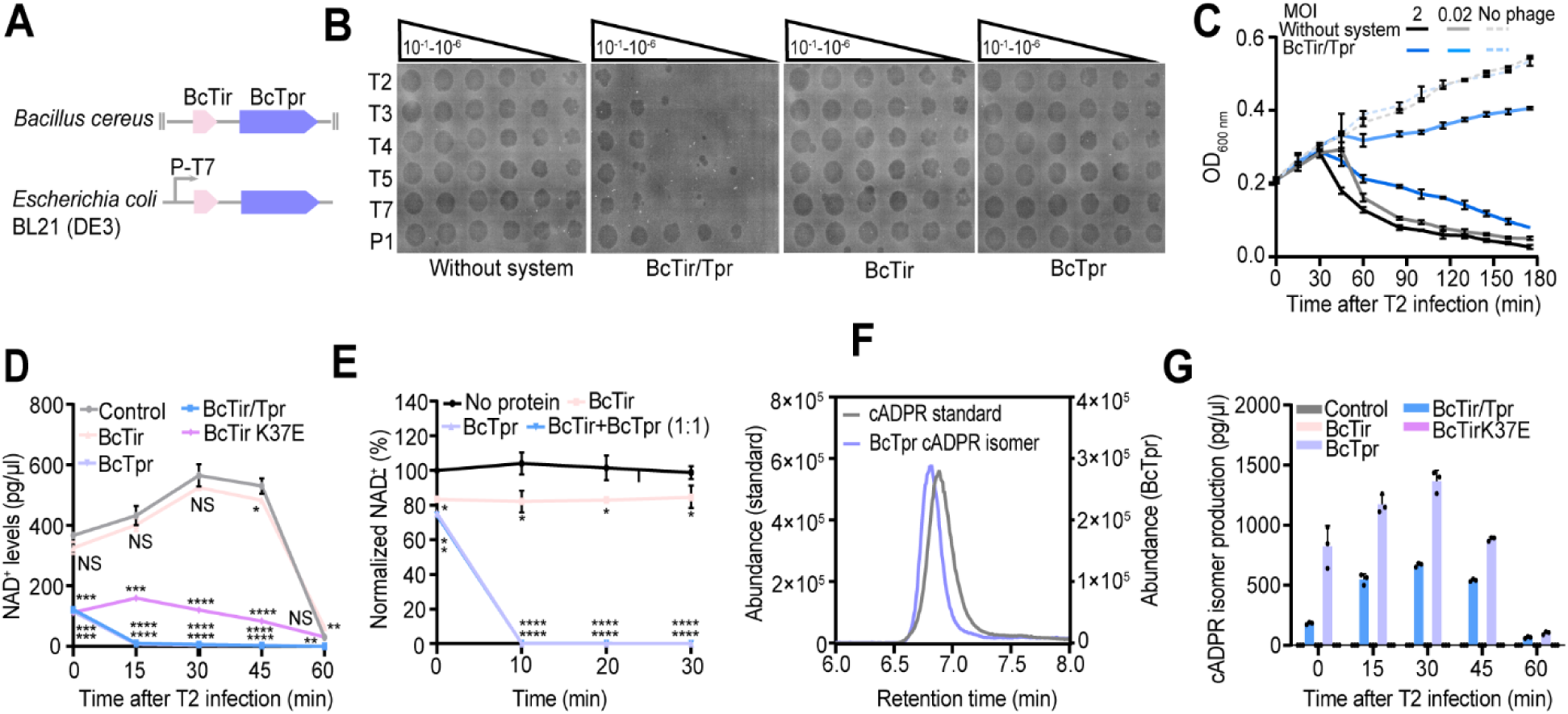
The BcTir/Tpr system exhibited anti-phage immunity via NAD^+^ metabolism. (A) Schematic graph of the BcTir/Tpr system in the genome of *B. cereus* which was cloned into the pET28a expression plasmid under the T7 promoter (P-T7) and then transformed into *E. coli* BL21 (DE3). In comparison, an empty pET28a vector was also transformed serving as a negative control. (B) Phage titrations in the cultures after infection. Culture supernatants in serial dilution were spotted onto LB plates bearing the BcTir/Tpr system, BcTir, and BcTpr, respectively, at 5 h post infection. T series are the lytic phages, and P1 is the lysogenic phage. (C) Growth curves in liquid cultures for *E. coli* containing the BcTir/Tpr system and *E. coli* transformant lacking the system (empty vector) were infected by T2 phage at 37°C. Three independent replicates were used for calculating the mean values and error bars represent the standard error of the mean (SEM). (D) Intracellular NAD^+^ degradation of various *E. coli* transformants measured by HPLC-MS at 15, 30, 45, and 60 min after phage infection. Time 0 represents uninfected cells. The “control” is *E. coli* without any parts of the BcTir/Tpr system. The other conditions are with the BcTir/Tpr system, BcTir, BcTir K37E, and BcTpr. (E) NAD^+^ depletion measurement of purified BcTir, BcTpr and BcTir + BcTpr (1:1 in molar ratio) (normalized to control at 0 min). Sample without proteins served as the negative control. (F) cADPR standard curve in grey (left Y-axis), and the cADPR isomer curve in light blue (right Y-axis) generated in BcTpr-containing cell lysates were identified by HPLC chromatograms. (G) cADPR isomer production measurements in different lysates, including control cells and cells containing BcTir/Tpr, BcTir, BcTir K37E, and BcTpr. The data from C, D, E, and G were shown as mean ± standard error of the mean (SEM). Data from D and E were compared to the (D) “Control” and (E) “No protein” conditions at the same time point using a two-tailed student’s *t*-test. “****” indicates *p-*value < 0.0001, “***” indicates *p-*value < 0.001, “**” indicates *p-*value < 0.01, and “*” indicates *p*-value < 0.05. “NS” means no significance.

### NAD^+^ depletion by BcTpr requires cADPR isomer produced at its N terminus

As we demonstrated above, BcTpr had dual functions, cADPR isomer production and NAD^+^ depletion, which have not been reported previously. Therefore, specifying the role of functional domains was investigated. Structural modeling of BcTpr using the AlphaFold program (Jumper et al., 2021) identified that its N terminus (amino acids: 1-189) (Figure 2A), BcTprN, resembles N-glycosidase, MilB, of *Streptomyces rimofaciens* (Zhao et al., 2014), suggesting that its glutamate residue is the core catalytic site responsible for the N-glycosidase activity (Sikowitz et al., 2013; Zhao et al., 2014). Interestingly, the MilB is the structural basis for the identification and recognition of the bacterial TIR domain (Essuman et al., 2018). Moreover, superimposition of the structure of BcTprN onto MilB (PDB: 4JEM) in complex with cytidine 5’-monophosphate (CMP) (Sikowitz et al., 2013; Zhao et al., 2014), revealed six potential residues of BcTprN, I10, M11, N80, P81, N82, and E86, and one residue, D217, at BcTprC that were computationally able to form a hydrogen bond with the ribose-derived substrates (Figure 2B). The E86 and I10 form a hydrogen bond with hydroxyl (2’- and/or 3’-OH) group of ribose respectively, and D217 with the amine (-NH_2_) group of cytosine. N80, P81, and N82 form a hydrogen bond with 5’-phosphate (Figure 2B). It is critically important the BcTpr E86 forms two potential hydrogen bonds and corresponds to MilB E103 in structure, which is the most likely catalytic site for cutting the N-glycosidic bond.

**Figure 2.**
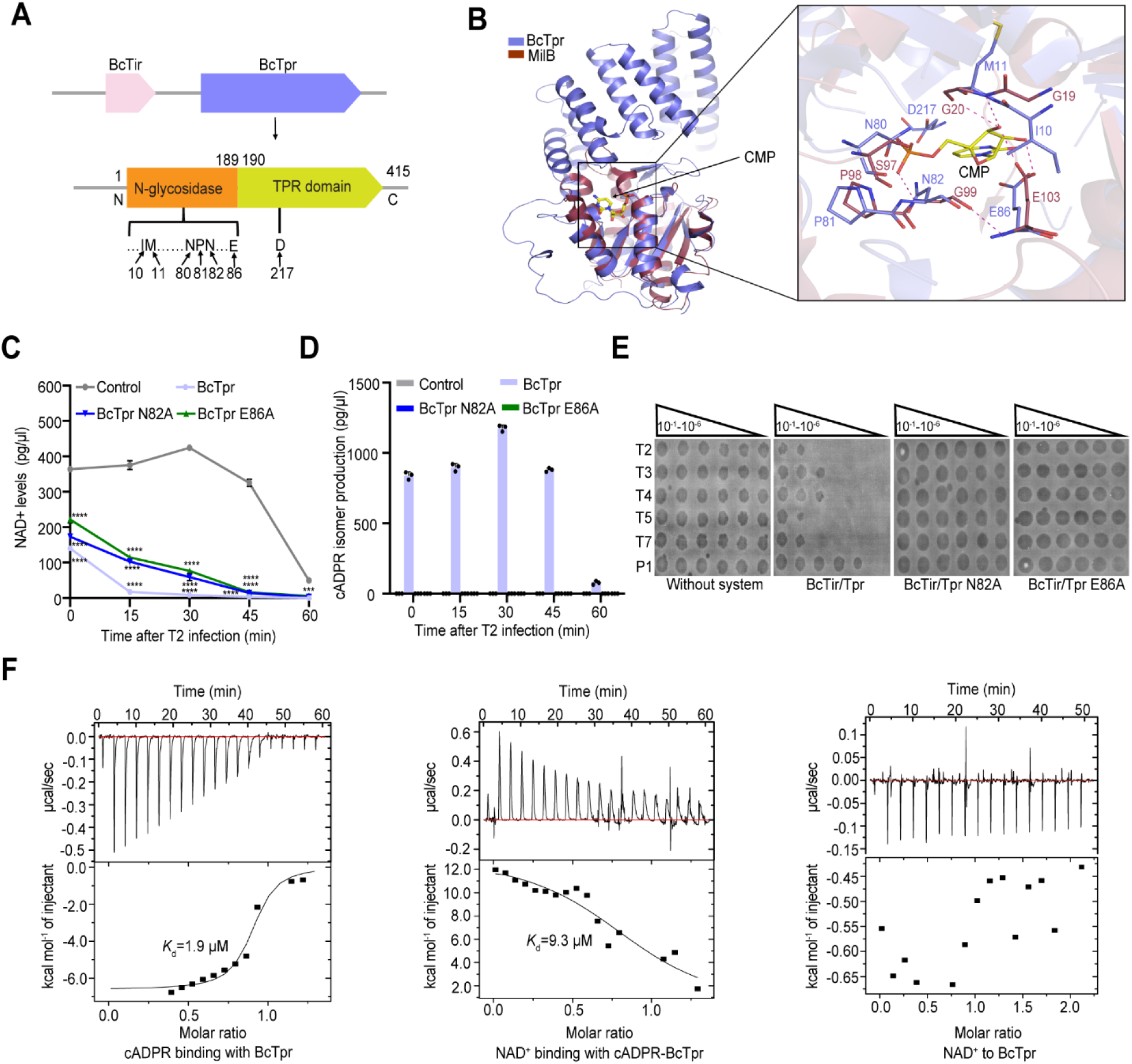
The cADPR isomer produced at the glycosidase domain of BcTpr, was essential for BcTpr to deplete NAD^+^. (A) Scheme of the domains and potential catalytic residues of BcTpr. BcTprN (amino acids: 1-189) was annotated as glycosidase; BcTprC (190-415) as the TPR domain. The labeled sites are potential catalytic residues. All the predicted residues mentioned above were mutated into alanine. (B) Superimposition of simulated BcTprN to MilB (PDB: 4JEM) formed a complex with cytidine 5’-monophosphate (CMP). The dash lines indicate potential hydrogen bonds. (C) Time points of intracellular NAD^+^ remaining in the lysates expressing BcTpr, BcTpr N82A, and BcTpr E86A. Lines represent the mean of three experiments. (D) Time points of cADPR isomer production in the lysates expressing BcTpr, BcTpr N82A and BcTpr E86A. Bars indicate the average of three experiments with the individual points overlaid. (E) BcTir/Tpr N82A and BcTir/Tpr E86A mutations abolished the antiphage immunity of the BcTir/Tpr system. Tenfold serial dilution plaque assays with T2 phage are displayed. (F) BcTpr binds to cADPR and the resulting complex allows further NAD^+^ binding. Isothermal Titration Calorimetry (ITC) detected the interaction between the cADPR and BcTpr. The cADPR-BcTpr complex can further bind NAD^+^. *K*d, dissociation constant. The data from (C) and (D) are shown as mean ± SEM. Data from C was analyzed by comparing to the “Control” group at the same time point by using a two-tailed student’s *t*-test. “****” indicates *p-*value < 0.0001, “***” indicates *p-*value < 0.001, “**” indicates *p-*value < 0.01, and “*” indicates *p*-value < 0.05. “NS” means no significance.

Site-directed mutagenesis revealed that BcTpr N82A and BcTpr E86A mutations impaired NAD^+^ depletion (Figure 2C) and cADPR isomer formation (Figure 2D) of BcTpr from the start to 30 min before cell destruction after phage infection. Both mutated residues abolished the anti-phage immunity of the BcTir/Tpr system (Figure 2E) compared to the other residues (Figure S6). Taken together, these two sites, BcTpr N82, and BcTpr E86, are the critical catalytic sites for executing the NAD^+^-depleting activity of the BcTir/Tpr system to defend against phage infection. Noticeably, NAD^+^ depletion was positively associated with the production of the cADPR isomer. Therefore, we further illuminated the relationship between BcTpr, NAD^+^, and the cADPR isomer and noted that cADPR as the analog replacing cADPR isomer. As shown in Figure 2F, cADPR was closely bound to BcTpr with a *K*d = 1.9 µM, which further promoted access of NAD^+^ to the BcTpr-cADPR complex with a *K*d = 9.3 µM. However, NAD^+^ did not show significant interaction with BcTpr in the absence of cADPR. Thus, cADPR isomer production is a prerequisite for the NAD^+^-depleting activity of BcTpr, suggesting an autoinhibitory configuration of BcTpr without phage infection akin to human SARM1 (Shi et al., 2022).

### The TPR domain of BcTpr regulates production of the cADPR isomer

Although we had identified that the glycosidase domain of BcTpr is responsible for producing the cADPR isomer, the role of a truncated N terminus and C terminus was still undefined. Therefore, truncation of the functional domains of BcTpr was carried out. As revealed in Figure 2A, BcTprN functioned as the glycosidase and bioinformatic analysis revealed that BcTprC (amino acids: 190-415) bore high structural similarity to the TPR domain (Figure 3A), which in most cases, is reported to act as a co-chaperone to modulate the enzymatic activities of downstream effectors (D’Andrea and Regan, 2003; Prodromou et al., 1999; Scheufler et al., 2000), such as Hsp90 ATPase. In addition, the TPR domain can confer antiviral defense (Abbas et al., 2017; Abbas et al., 2013) and direct glycosidic bond recognition and selection to guide the enzymatic activity for post-translational modifications (Allan and Ratajczak, 2011; Ramirez et al., 2021). It is reasonable to assume that the versatile functions of the TPR domain make it widely distributed to mediate various biological processes.

**Figure 3.**
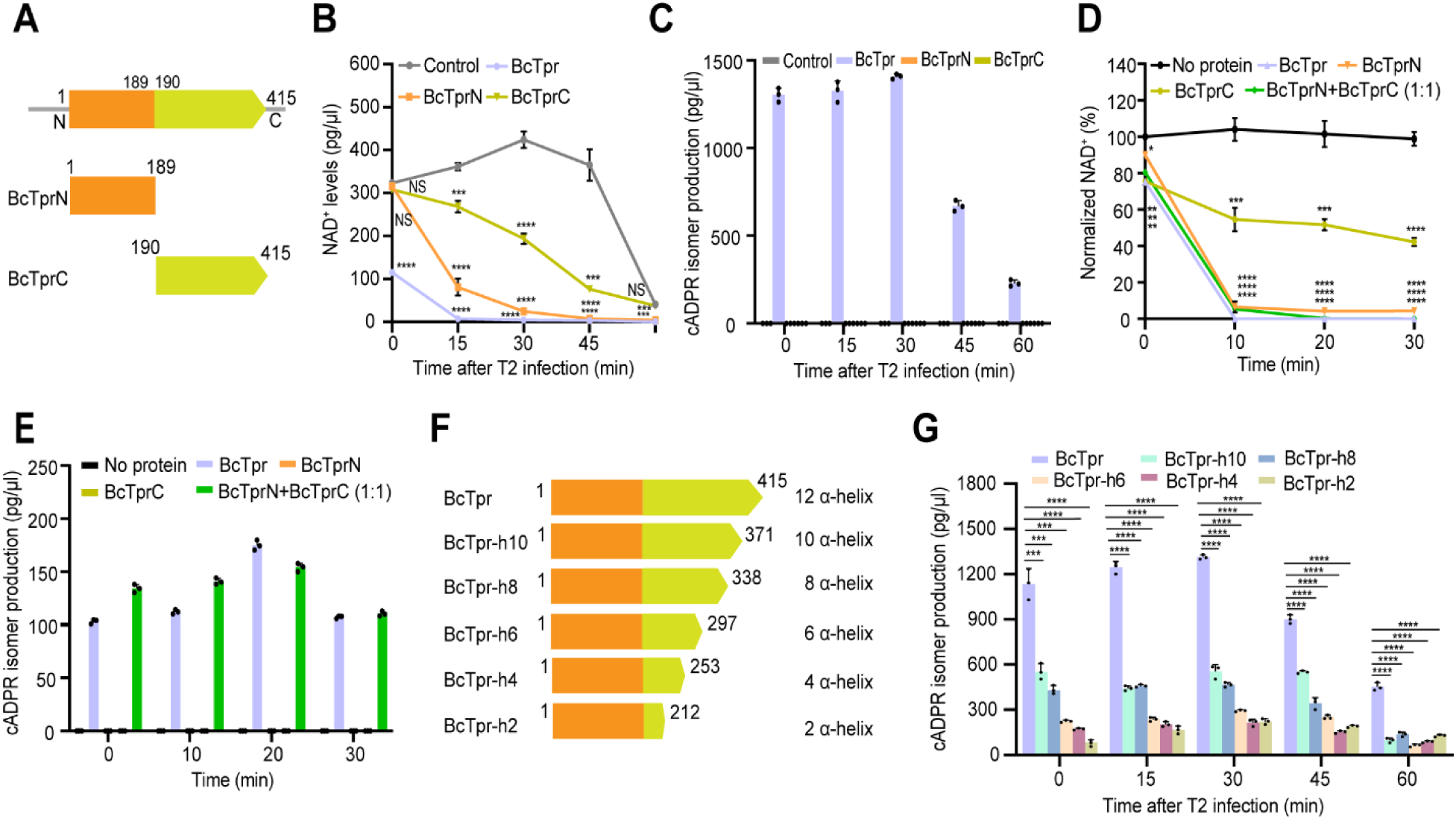
BcTprC guided the BcTprN on specific N-glycosidic bond cleavage of NAD^+^. (A) Two functional domains of BcTpr (amino acids: 1-415), BcTprN (1-189) and BcTprC (190-415). (B) Cellular NAD^+^ degradation of various *E. coli* transformants measured by HPLC-MS at indicated time points, 0 min, 15 min, 30 min, 45 min and 60 min, after phage infection. The “control” represents the *E. coli* without any parts of the BcTir/BcTpr system. Other groups are BcTpr, truncated N-terminus BcTprN, and truncated C-terminus BcTprC. (C) Production of the cADPR isomer was quantified at each time point from the lysates expressing control, BcTpr, BcTprN and BcTprC, respectively. (D) NAD^+^ depletion measurement of purified BcTpr, BcTprN, BcTprC and mixture of BcTprN + BcTprC in 1:1 molar ratio (normalized to control at 0 min). Samples without protein served as the control, No protein. (E) Production of the cADPR isomer was measured of BcTpr, BcTprN, BcTprC and mixture of BcTprN + BcTprC in 1:1 molar ratio (normalized to control at 0 min). Samples without protein served as the control, No protein. (F) Truncated BcTpr derivatives were generated by decreasing helix numbers of the TPR domain, BcTpr (12 helix), BcTpr-h10 (10 helix), BcTpr-h8 (8 helix), BcTpr-h6 (6 helix), BcTpr-h4 (4 helix) and BcTpr-h2 (2 helix). (G) Production of the cADPR isomer was quantified at each time point in the lysates expressing BcTpr, BcTpr-h10, BcTpr-h8, BcTpr-h6, BcTpr-h4, and BcTpr-h2. Bars indicating the average of three experiments with the individual points overlaid are shown. (B), (C), (D) (E) and (G) were obtained from three independent replicates. The data are presented as mean ± SEM. Data from B, C, D, E and G were analyzed compared to the “Control” (B, C), “No protein” (D, E) and BcTpr (G) by using two-tailed student’s *t*-test. “****” indicates *p-*value < 0.0001, “***” indicates *p-*value < 0.001 and “**” indicates *p-*value < 0.01, “*” indicates *p*-value < 0.05. “NS” means no significance.

Our biochemical results revealed the ability of BcTprN to deplete NAD^+^ *in vivo* (Figure 3B) and *in vitro* (Figure 3D), which agreed with the characterization in Figure 2, but with less efficiency compared to the intact BcTpr. The incubation of BcTprN to BcTprC (1:1, molar ratio) can restore the NAD^+^-depleting ability as BcTpr in Figure 3D. Intriguingly, BcTprC appeared to deplete NAD^+^ partially *in vivo* (Figure 3B), which could be the result of TPR-mediated indigenous enzymatic activity of NAD^+^ hydrolysis as less NAD^+^ depletion was noticed *in vitro* (Figure 3D). Of more interest, cADPR isomer was not produced by the separated BcTprN and BcTprC *in vivo* (Figure 3C) and produced when both incubated at 1:1 molar ratio (Figure 3E), which indicated that BcTprN was directed by the TPR domain, BcTprC, to target the glycosidic bond of NAD^+^ for generating the cADPR isomer. Moreover, the fact that the released or sole BcTprN depleted NAD^+^ suggested a prohibitive action of BcTprC on BcTprN in the integrated BcTpr, which required the cADPR isomer to allow access to NAD^+^ and its further depletion as shown in Figure 2F. Further, BcTpr derivatives were constructed by decreasing the number of helices in the TPR domain from 12 helices as shown in Figure 3F. It was evident that cADPR isomer production by truncated derivatives of BcTpr was significantly reduced indicating that less helices in the TPR domain of BcTpr results in less cADPR isomer production (Figure 3G). These results agreed with the observation of truncated BcTprN without any helices in the TPR domain in which cADPR isomer was not produced. Interestingly, every four helix decrease of the TPR domain starting from BcTpr to BcTpr-h8 and BcTpr-h4, halved cADPR isomer production under phage infection excluding the 60 min time point which suggested that four helices could represent a critical functional unit of the TPR domain. Altogether, the TPR domain was required for the production of the cADPR isomer by the glycosidase domain, BcTprN, of BcTpr.

### Pleiotropic roles of the BcTir/Tpr system against viral infection

In TPR-involved anti-viral cases, such as the innate immune effector IFITs, viral proliferation is inhibited by prohibiting the translation of viral mRNA (Abbas et al., 2017). Since bacterial growth arrest was delayed (MOI = 2) and abolished (MOI = 0.02) by the BcTir/Tpr system (Figure 1C), phage replication was accordingly assumed to be disturbed. We utilized quantitative PCR (qPCR) with phage-specific sequence to detect the adsorption and DNA replication of T2 phages in the recipient cells. We collected the supernatant samples after centrifugation for the adsorption assay at 30 min prior to the cessation of cell growth and precipitated samples for phage replication within one hour after phage infection. Importantly, the supernatants of cells with the BcTir/Tpr system exhibited significantly higher copy numbers of phage genome than those without the system prior to or at 20 min after infection revealed phage adsorption impairment by the BcTir/Tpr system (Figure 4A), which might be alteration of membrane recognition (Huiting and Bondy-Denomy, 2023). Furthermore, the internalized phages started replication after 15 min and the increase of phage copies was depressed in cells with the BcTir/Tpr system approximately five-fold compared to those without before cell destruction at 30 min (Figure 4B). Although more control cells were lysed at 60 min compared to those with the BcTir/Tpr system, the precipitated control cells also contained over five-fold times more phages than cells with the BcTir/Tpr system. By contrast, separated BcTir or BcTpr could not impede phage infection (Figure S7). Moreover, transmission electron microscopy (TEM) results confirmed that the BcTir/Tpr system impaired phage proliferation and prevented the subsequent cell lysis (Figure S8). It was evident that the intact BcTir/Tpr system was essential for anti-viral immunity. Since both TIR and TPR have been reported to have protein-protein interaction potential, this was detected by pull-down experiments (Figure 4C) and Bio-Layer Interferometry (BLI) (Figure S9A). In addition, BcTir was capable of binding with BcTprN and BcTprC separately (Figure S9B-E).

**Figure 4.**
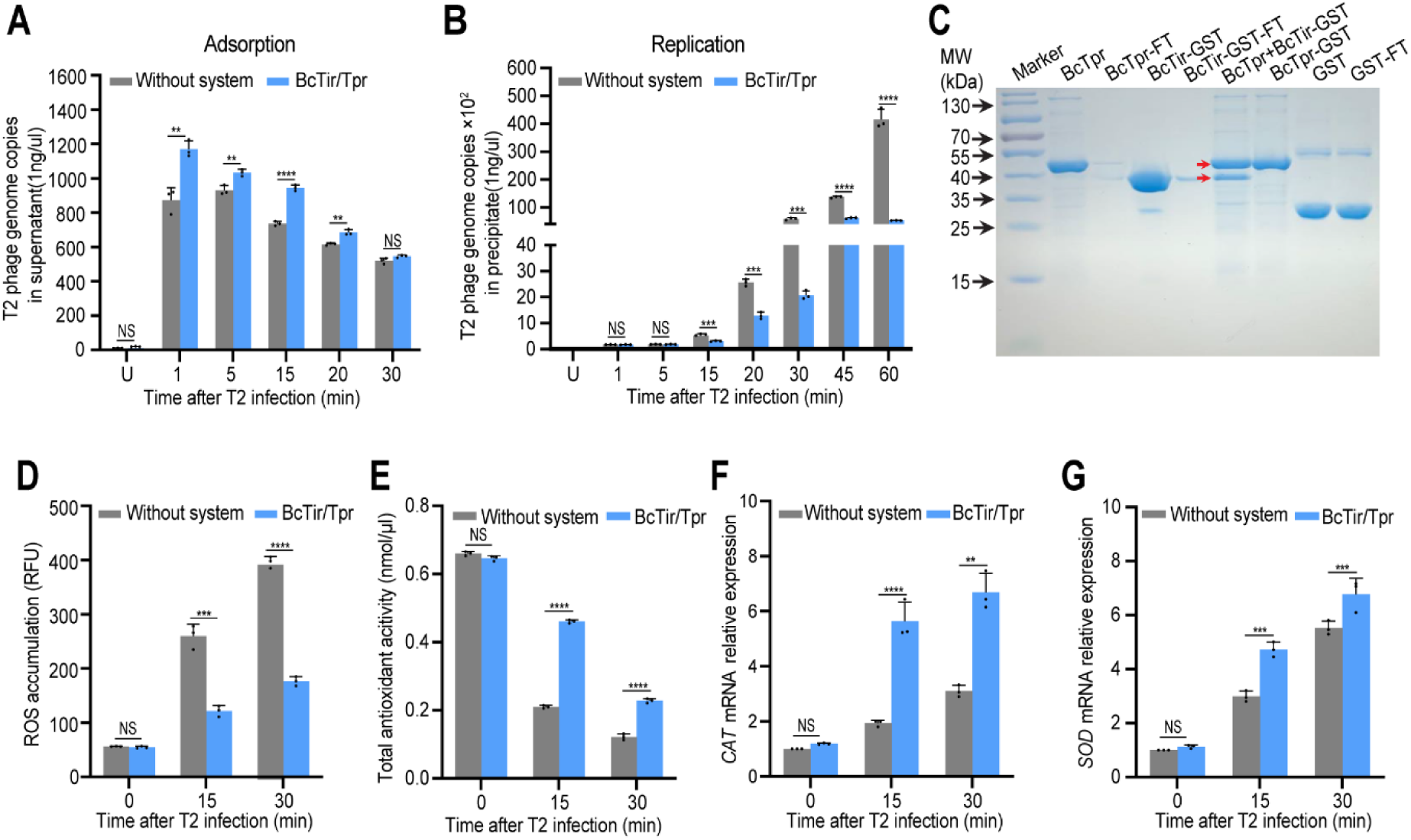
The BcTir/Tpr system prevented phage proliferation and protected cell viability from ROS oxidation induced by phage infection. (A) Phage adsorption was assessed by detecting the copy numbers of phage in the supernatant via qPCR following a time courses during 30 min phage infection. More copies of phage genomes in the supernatant indicated weaker phage adsorption. (B) Replication of phage genomes measured by qPCR in the precipitated population during the time course mentioned above. (C) The interaction between BcTir and BcTpr was observed by pull-down experiments. BcTir was tagged with GST, which was used as the negative control to rule out non-specific interaction mediated by the GST tag as well as no interaction between His tag and BcTir (data not shown). Marker: protein ladder of different molecular weights; FT: flow through. The upper arrow represents BcTpr and lower is BcTir-GST. (D)-(G) Cellular ROS accumulation (D), total antioxidant activity (E), relative expression at the mRNA level of the catalase gene (F) and superoxide dismutase gene (G) of cells with or without the BcTir/Tpr system were measured at 0 min, 15 min, and 30 min after T2 phage infection. “U” in (A) and (B) indicates uninfection. Bars represent the average values of three independent experiments with individual data points shown. Data is presented as mean ± SEM. Data from (A), (B), and (D)-(G) were analyzed compared to the “Without system” at the same time point by using two-tailed student’s *t*-test. “****” indicates *p-*value < 0.0001, “***” indicates *p-*value < 0.001, “**” indicates *p-*value < 0.01, and “*” indicates *p*-value < 0.05. “NS” means no significance.

Reactive oxygen species, ROS, are an important signal for many biological processes but their dramatic high-level generation following phage entry through the host membrane generates detrimental effects such as protein oxidation and cell death (Dong et al., 2015; Johnson et al., 2022; Liu et al., 2021). In general, organisms take immediate measures to minimize the damage of excessive ROS including upregulation of scavenging enzymes. In our work, we found that intracellular ROS were highly induced in the non-BcTir/Tpr system, BcTir and BcTpr, in which ROS levels were over three times than that of the BcTir/Tpr system at 15 and 30 min after phage infection (Figure 4D and Figure S10A). Even though there was a loss of antioxidant activity of all cells upon phage infection, the BcTir/Tpr system remarkably salvaged up to 50% of activity following the time course compared to controls without the system, which were limited to 20% (Figure 4E), and BcTir and BcTpr alone (Figure S10B). Subsequently, separated BcTir and BcTpr were not taken into account for elaborating the underlying mechanism. We examined two well-studied ROS-removal enzymes, catalase (CAT) and superoxide dismutase (SOD). The CAT gene was upregulated approximately two fold in cells with the BcTir/Tpr system compared to the non-system counterpart. SOD gene expression was also markedly increased by the BcTir/Tpr system. Accordingly, the BcTir/Tpr system played a protective role during phage infection by upregulating the expression of ROS scavengers to maintain homeostasis of ROS.

### BcTpr as a core architecture is widely distributed

The above data sufficiently supported the anti-viral immunity role of the BcTir/Tpr system adopting multiple strategies to modulate NAD^+^ metabolism, viral proliferation, and ROS homeostasis (Figure 5A), in which BcTpr played dual roles of cADPR isomer production and NAD^+^ depletion. Thus, BcTpr was considered the core architecture of the BcTir/Tpr system as previously proposed (Burroughs and Aravind, 2020). To clearly illustrate the diversity of Tpr-based assemblies and glean insight into their potential function, we analyzed the prokaryotic assemblies from available databases (Burroughs and Aravind, 2020). We found that 78 operons contained the core Tpr architecture that were divided into seven groups according to the variance of TIR-containing neighboring genes from simplicity to complexity (Figure 5B). The core Tpr architectures alone were the primary findings (29/78), and were distributed widely and mainly in Proteobacteria phyla accounting for over 50%. Tir-Tpr assemblies ranked second (28/78) and included most of the strains from Firmicutes minus two, which only bore the core Tpr architecture, indicating that the Tir-Tpr assemblies are evolutionarily conserved in the Firmicutes phyla. Similar to the TIR domain, YspA also belonged to the SLOG super family and is capable of binding to nucleotides and ligands to regulate cell division (Brzozowski et al., 2019; Burroughs and Aravind, 2020). There were five YspA-Tpr assemblies identified, among which four belonged to Proteobacteria and one to Bacteroidota. In addition, Patatin from plants has been reported as holding multiple functions, such as oxidation resistance, lipid metabolism, and energy homeostasis (Liu et al., 2003; Rydel et al., 2003). Four Patatin-Tpr assemblies were discovered, in which three were from Proteobacteria and one from Bacteriodota. In addition, two different proteins were observed to accompany the core Tpr architecture individually, the LSDAT domain belonging to the SLOG superfamily, which also has nucleotide-binding potential (Burroughs et al., 2015), and the calcineurin domain that is assumed to function as a phosphodiesterase, capable of targeting cyclic nucleotides and modulating or terminating signal delivery (Burroughs and Aravind, 2020). Interestingly, the above-mentioned domains can be fused and rearranged together to accompany the core Tpr, which could be the result of evolutionary adaptation. Importantly, these domains have also been characterized to involve the eukaryotic immune response to invading pathogens. Therefore, further demonstration and examination of these Tpr-centered assemblies will add substantial understanding to both prokaryotic and eukaryotic immunity.

**Figure 5.**
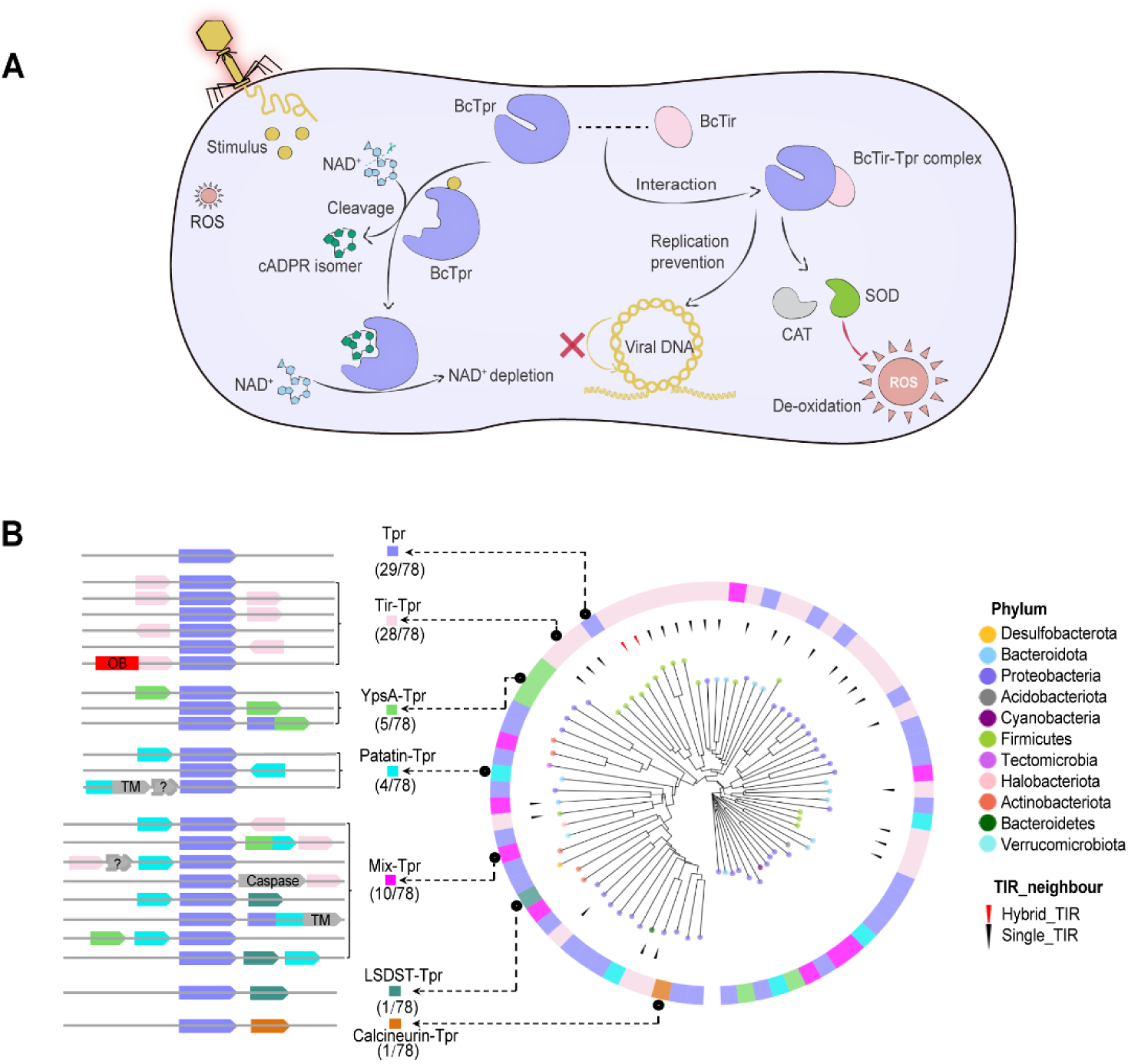
Proposed working mechanism of the BcTir/Tpr system and (Bc)Tpr-centered assemblies in represented prokaryotic genomes. (A) Upon T series phage infection, immediate stimuli are produced, which can bind and allosterically activate the BcTpr to enzymatically cleave NAD^+^ into the cADPR isomer, and the newly formed complex of cADPR isomer and BcTpr allows further NAD^+^ depletion. The BcTir/Tpr complex reduces viral proliferation through replication prevention. In addition, the complex upregulates CAT and SOD to scavenge excessive ROS and maintain oxidative homeostasis. De-oxidation: decrease oxidation. (B) Phylogenies and assemblies of 78 Tpr homologs across the prokaryotic phyla. Tips of the tree in different colors (solid circles) represent their taxonomic phyla. Arrows indicate that the core Tpr architecture that is accompanied by a gene containing the TIR domain only (black) and TIR domain as the subunit (red). The outer ring depicts the designated types of Tpr-centered assemblies. The operons only represent the order of component genes arrangement. “OB”: OB-fold nucleic acid binding domain (Theobald et al., 2003); “TM”: transmembrane domain; “?”: unidentified protein; “Caspase”: cysteine protease (Boatright and Salvesen, 2003).

## Discussion

Recently, many studies on bacterial innate immune systems have highlighted the importance of the broadly-distributed TIR domain, which catalyzes NAD^+^-derived signal production and triggers the immune response of the host to prevent invasion escalation (Essuman et al., 2022; Huang et al., 2022; Jia et al., 2022; Ofir et al., 2021). Protein assembly hybrids with the TIR domain diversify this signal-dependent immune system (Coronas-Serna et al., 2020; Huang et al., 2022; Jia et al., 2022; Koopal et al., 2022; Shi et al., 2022). Our work studied TIR-containing assemblies in tandem with a universal scaffold domain, TPR (D’Andrea and Regan, 2003; Yang et al., 2005; Zeytuni and Zarivach, 2012), and revealed their synergistic relationship for anti-phage defense in the representative operon, BcTir/Tpr system, from *Bacillus cereus*. Intriguingly, BcTpr, rather than BcTir, exhibited NADase activity producing the NAD^+^-based cADPR isomer which in turn forms a complex with BcTpr to allow the binding of NAD^+^ and escalates its depletion during phage infection. In addition, the N terminus of BcTpr bore dual roles for cADPR isomer formation directed by the TPR domain at the C terminus of BcTpr, and signal-dependent NAD^+^ depletion. Further, BcTir-BcTpr interaction aligned to the plaque titrations and was required for the blockade of phage proliferation and phage-induced ROS homeostasis.

The crystal structure of BcTir characterized its high structural similarity to the TIR domains of ThsB of the Theories system and human SARM1 (Figure S4A) (Eastman et al., 2022; Huang et al., 2022; Jia et al., 2022; Ofir et al., 2021; Tian et al., 2021) and was therefore assumed capable of generating NAD^+^-derived signals. Unexpectedly, the NADase activity of BcTir was inactive and might be due to a natural mutation of BcTir at the catalytic site from glutamate (E) to lysine (K). The BcTir K37E mutant maintained low-level NAD^+^ as well as the BcTir/Tpr system or BcTpr prior to phage infection and depleted NAD^+^ minimally without generating derived signals during phage infection. Clearly, BcTir played a noncanonical role in NAD^+^ metabolism. Further, BcTir was required in the BcTir/Tpr system for depression of viral proliferation (Figure 4B and Figure S7), supporting an indispensable role of BcTir for anti-viral immunity (Figure 1B). Protein-protein interaction revealed that the BcTir/Tpr system worked in coordination as a BcTir-BcTpr complex (Figure 4C). The N terminus (amino acids: 1-189) of BcTpr, resembles a glycosidase (Essuman et al., 2018; Ka et al., 2020) in structure (Figure 2B) and the site mutants, BcTpr N82A and BcTpr E86A, abolished cADPR isomer production and attenuated NAD^+^ degradation, which unraveled a dual role of the N terminus of BcTpr. Noticeably, the TPR domain at the C terminus of BcTpr was required for NAD^+^-derived signal production by the glycosidase domain of BcTpr at the N terminus that accelerated subsequent NAD^+^ depletion (Figure 2F). The truncated N terminus of BcTpr, BcTprN, also retained the ability to deplete cellular NAD^+^ (Figure 3B). As reported recently, the TPR domain can guide the glycosite selection of the conjugated enzymatic domain for post-translational modification (Ramirez et al., 2021). We are the first to report that the TPR domain directs glycosidase to target an N-glycosidic bond producing a cADPR isomer and the amount of signal production is regulated by the alpha-helix number in the TPR domain of BcTpr (Figure 3F and G). Further study of critical residues within the alpha helix of the TPR domain would significantly increase understanding of this helix regulation. For example, the enzymatic domain assembled with TPR could be repurposed with glycosite-related modifications, such as O-linked N-acetylglucosamine (O-GlcNAc) installation (Allan and Ratajczak, 2011; Ramirez et al., 2021).

In the well-studied TPR-containing enzymes, the TPR domain acts in a co-chaperone manner to prohibit enzymatic activity under normal conditions and switches the enzyme to an active state via binding the stimulators produced under changing or stressful environments, such as viral infection (Abbas et al., 2017; Abbas et al., 2013). Thus, BcTpr, especially its N terminus, is likely to be inhibited by its TPR domain prior to phage infection. Phage attack produced immediate ROS production (Figure 4D) as an emergent signal to activate downstream effectors, such as the TPR domain following a chaperone pattern (D’Autréaux and Toledano, 2007). Active BcTpr in physiological conditions produced the cADPR isomer at its N terminus and further accelerated NAD^+^ depletion (Figure 5A). Intriguingly, the BcTir/Tpr system also worked in a chaperone-like fashion to protect antioxidant activity by upregulating expression of ROS scavengers (Figure 4E-G), which indicated a critical role for the TPR domain. Furthermore, the BcTir/Tpr system significantly impaired the invading phage proliferation (Figure 4A and B), which could be the consequence of TPR domain activity deterring viral mRNA translation preferentially via formation of an RNA-binding tunnel as described for human IFITI (Abbas et al., 2017; Abbas et al., 2013). In addition to the TPR domain, the N terminus of BcTpr functioned as glycosidase which recognized 5’-hydroxymethylcytosine (5hmC) that was incorporated into the phage genome as an epigenetic modification during replication (Song et al., 1999). In general, BcTpr appears to wrap up the phage DNA-mRNA complex to impede viral transcription and translation in agreement with the phage replication results (Figure 4B). Nevertheless, these assumptions await further investigation. We characterized the role of our BcTir/Tpr system, especially BcTpr, as the core architecture (Figure 5B) which was widely distributed and assembled with multiple immune effector domains. This observation indicated a ubiquitous function of BcTpr for innate immune systems.

Collectively, the reported BcTir/Tpr system was responsible for an NAD^+^-derived signal-dependent anti-viral immune mechanism. The defense was executed in a salvage manner, similar to the abortive infection of the ThsA/ThsB system and delayed death of infected cells likely through a depression of phage proliferation and maintenance of ROS homeostasis. Due to the prerequisite of both BcTir and BcTpr and its highly intriguing coordination when facing phage invasion, some of our intriguing observations are distinct from previously discovered mechanisms and our studies may richly increase the knowledge of anti-phage immunity in bacteria. Altogether, anti-viral immunity conferred by the BcTir/Tpr system is multifaceted and interconnected in an intricate nature which demands continued research.

## Acknowledgements

We are grateful to Bryon Grove and Laurie Kim Young at the Image Core of UND for the assistance in TEM. We thank Steven Adkins and Suba Nookala for the assistance in FACS at the Flow Cytometry Core of UND. LC-MS/MS analysis was performed at the UND SOMHS MS core facility supported by the SOMHS Dean’s office. This project was supported by the National institute of Health for Grants R01 AI1138203 for M.W. Flow Cytometry and Imaging Core services were supported by NIH/NIGMS award P20GM113123, DaCCoTA CTR NIH grant U54GM128729, and UND SMHS funds.

## Author contributions

Experiments were designed and performed by Y.L. and C.Z. Gene identification and phylogenetic analyses were performed by C.Z. and Y.L. M.G developed and performed the LC-MS/MS assay and associated data analysis. S.G developed and performed the LC-MS/MS assay and sample preparation method. N.K, J.S and C.Z. performed the structural analysis and ITC experiments. J.W and C.Z. drew the model of BcTir/Tpr system. Y.L., L.Z and A.G. performed the BLI experiments. C.C. and M.W. (initially conceived) supervised the study and wrote the paper together with Y.L. and C.Z.

## STAR★METHODS

### KEY RESOURCE TABLE

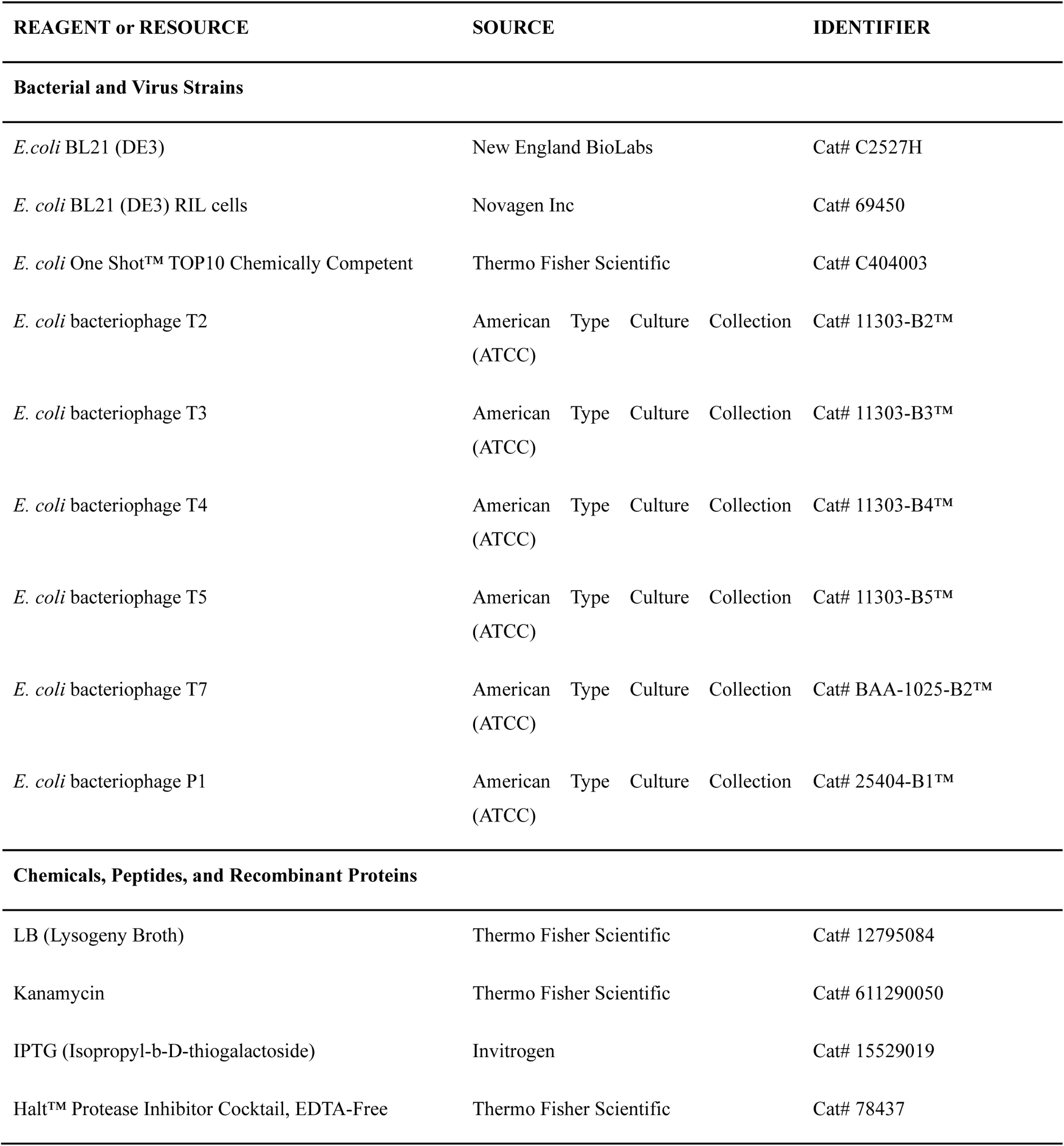
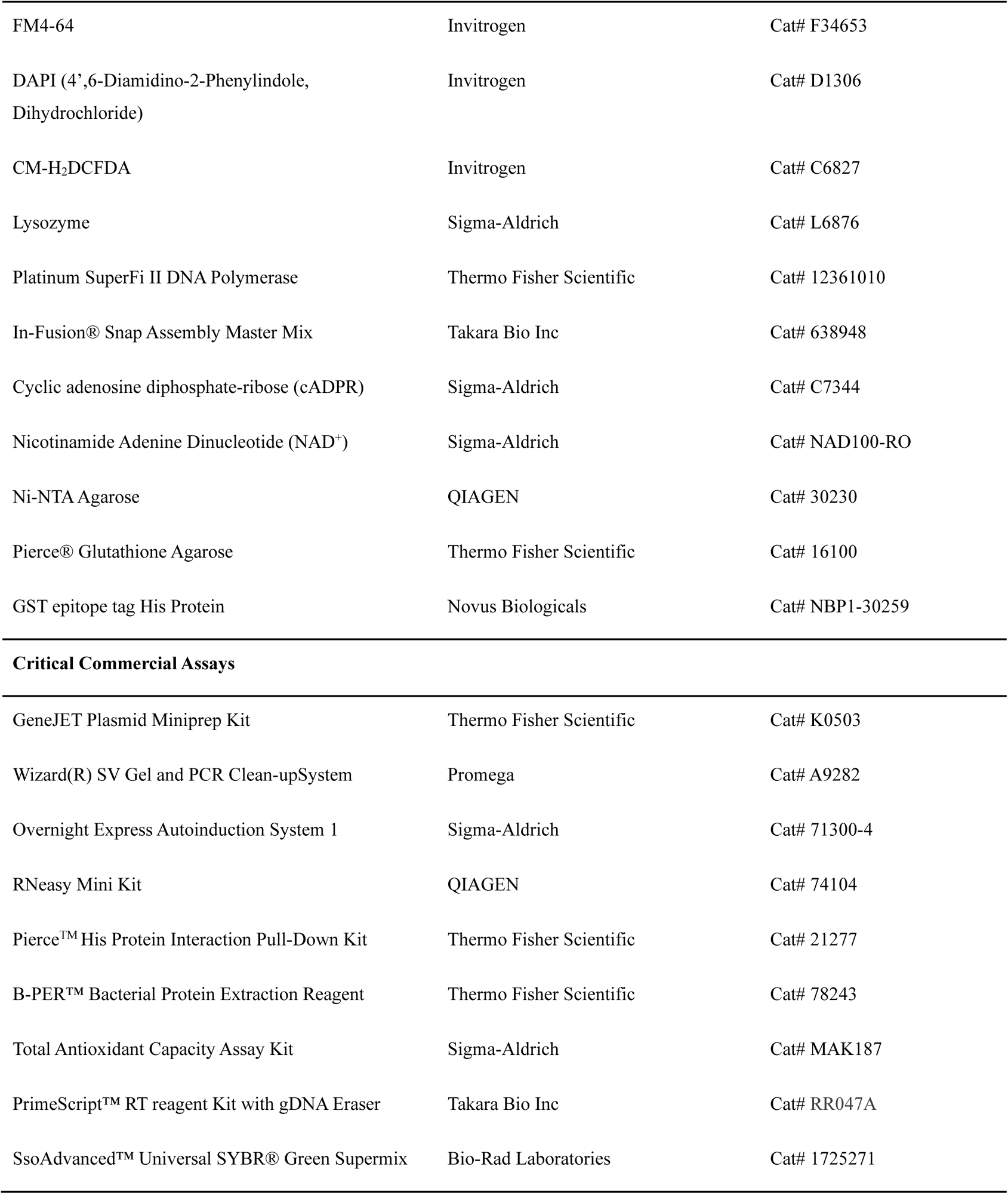
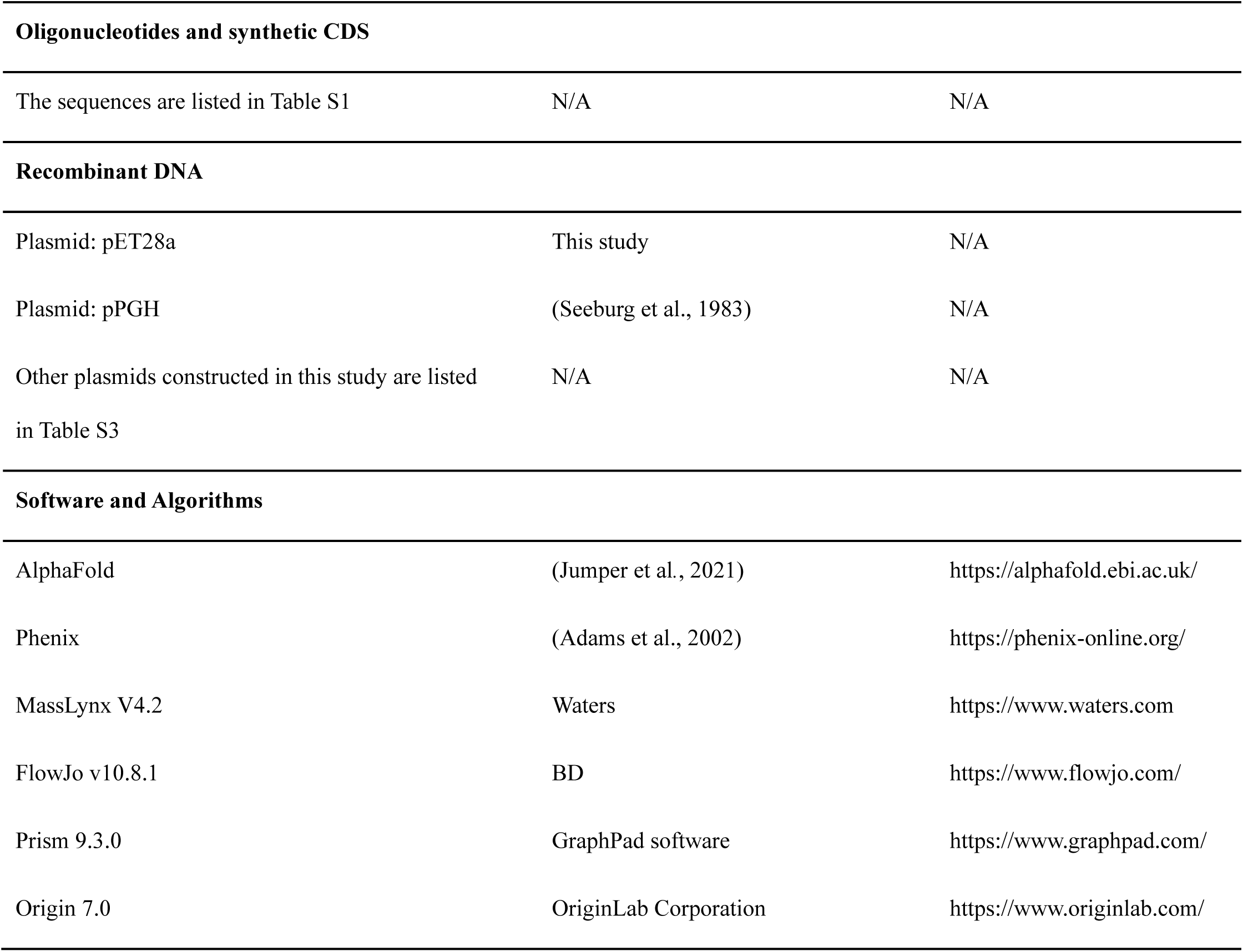

### RESOURCE AVAILABILITY

#### Lead contact

Further information and requests for resources should be directed to the lead contact, Colin Combs.

#### Materials availability

Plasmids, strains, virus and other unique reagents generated in this study are available upon request.

#### Data and code availability

Crystal structure of BcTir was deposited at the protein data base under identifier PDB: 8FZ9, and is publicly available as of the date of publication. All additional data reported in this paper will be shared by the lead contact upon request. This paper does not report original code.

### EXPERIMENTAL MODELS AND SUBJECT DETAILS

#### Bacterial strains growth conditions

*Escherichia coli* One Shot^TM^ TOP10 (Thermo Fisher Scientific, C404003) and BL21 (DE3) were routinely grown at 37°C with shaking at 220 rpm in LB medium or LB-agar medium supplemented with kanamycin (50 μg mL^-1^) when required and used for plasmid cloning and protein expression respectively.

### METHOD DETAILS

#### Plasmid and strain construction

All synthesized sequences are listed in Table S1. Plasmids used in the study are listed in Table S3. The full-length two-gene *Bctir* and *Bctpr* operon with 400 bp interregional upstream sequence of *Bacillus cereus* were synthesized and constructed into pET28a by Gene Universal Inc (Newark DE, USA). The *Bctir* with 400 bp interregional upstream sequence was amplified by PCR using the pET28a-BcTir/Tpr plasmid as the template, and cloned into pET28a vector. To generate the *Bctpr*, the 400 bp interregional sequence hybrid with *Bctpr* (containing 73 bp intergenic region) were fused using in-fusion snap assembly master mix kit, and cloned into pET28a vector. A similar strategy was used to construct the BcTir and BcTpr variants. For the site-directed mutagenesis of BcTpr with the interregional and intergenic sequence (ints), the amplified left arm contains the ints and the site-mutated regions, which was overlapped with the amplified right arm. The generated pET28a-BcTpr was the template for the above amplification. Both arms were fused into a single fragment and cloned into pET28a. For the BcTir/BcTpr system with the site mutant, the left arm is the BcTir with upstream interregional sequence and the right arm is the site-mutated BcTpr with an intergenic sequence. All the left arms used the pET28a-BcTir/Tpr as the template, and all the right arms used the site mutants of pET28a-BcTpr as the template. Both arms were fused into a single fragment and cloned into pET28a. The amino acids, I10, M11, N80, P81, N82, E86 and D217 of BcTpr were mutated into alanine. Strategy to construct BcTpr-h10 to BcTpr-h2 was similar to the above. All the primers are listed into Table S2.

For separate expression of *Bctir* and *Bctpr*, the interregional sequence and intergenic region were excluded. The truncated proteins BcTprN and BcTprC followed the same method. All recombinant plasmids and the empty pET28a vector were first transformed into *E. coli* One Shot™ TOP10 and then introduced into the *E. coli* host strain BL21 (DE3).

#### Phage cultivation

*E. coli* phages T2, T3, T4, T5, T7, and P1 were propagated by infection of *E. coli* BL21 at 37°C with 220 rpm shaking until the cells were lysed completely. The lysates were centrifuged and the supernatants were filter sterilized through a 0.22-μm filter and stored in SM buffer (100 mM NaCl, 8 mM Mg_2_SO_4_, 50 mM Tris-HCl, pH 7.5) over chloroform at 4°C.

#### Plaque assays

Phage titer was determined using the small drop plaque assay method as previously described (Mazzocco et al., 2009). In brief, 300 μL of an overnight bacterial culture were mixed with 10 mL of 0.7% LB agar medium supplemented with 10 mM Mg_2_SO_4_ and 50 μM IPTG in plates. After solidification, 3 μL of 10-fold serial dilutions of phage lysate were spotted onto the surface. Plates were incubated overnight at 37°C. Plaque forming units (PFUs) were determined by counting the derived plaques after overnight incubation and lysate titer was determined by calculating PFUs per mL.

Plaque assays were performed as previously described with some modifications (Mazzocco et al., 2009). *E. coli* BL21 (DE3) strains carrying recombinant plasmids BcTir, BcTpr, and negative control pET28a, were grown overnight at 37°C. Then, 300 μL of an overnight bacterial culture was mixed with 10 mL of 0.7% LB agar medium supplemented with 10 mM Mg_2_SO_4_ and 50 μM IPTG in plates. After solidification, 3 μL of 10-fold serial dilutions of phage lysate were spotted onto the surface. Plates were incubated for 5 h at 37°C.

#### Phage-infection dynamics in liquid medium

Overnight bacterial cultures were subcultured 1:100 into 12 mL of fresh LB medium supplemented with 50 μg mL^−1^ kanamycin and 50 μM IPTG and grown with shaking at 37°C until the optical density at 600 nm (OD _600_) reached 0.4. Phage T2 was added at a multiplicity of infection (MOI) of 0.02 or 2. Uninfected cultures were used as controls. 200 μL samples were removed every 15 minutes and transferred into a 96-well plate. The plate was read in a BioTek Synergy HTX Plate Reader Multi-Mode Microplate Reader (Bio-Tek, Winooski, VT, USA).

#### Protein expression and purification of BcTIR and BcTPR for biochemical and structural characterizations

For biochemical characterization, the purification procedure for all the proteins and related mutants were similar except for BcTprN. Since the expression of BcTprN was very low, we used the overnight express autoinduction system 1 to induce the expression of BcTprN without adding isopropyl β-d-1-thiogalactopyranoside (IPTG). Overnight cultures of *E. coli* BL21 (DE3) carrying the indicated plasmids were subcultured 1:100 and grown at 37°C to an OD _600_ of 0.6, then induced with 0.1 mM IPTG for 15 hours at 16°C. The cells were harvested by centrifugation at 5000 g for 10 min and the pelleted cells were suspended in 30 mL of lysis buffer (50 mM sodium phosphate, pH 7.4, and 300 mM NaCl, 10% glycerol). Cells were lysed by sonication on ice and then centrifuged (16,000 g at 4°C for 50 min). The cell extract was loaded onto Ni-NTA agarose resin. The background proteins were washed away with lysis buffer containing 40 mM imidazole. After washing, the proteins were eluted with an elution buffer (50 mM sodium phosphate, pH 7.4, 300 mM NaCl and 200 mM imidazole). His-tagged proteins were eluted using a lysis buffer containing 250 mM imidazole. The eluent was concentrated using a 3 kDa ultra-centrifugal filter (Sigma-Aldrich, UFC900324) and dialyzed overnight at 4°C in lysis buffer (50 mM sodium phosphate, pH 7.4, and 300 mM NaCl, 10% glycerol).

For structural characterization, *E. coli* BL21 (DE3) RIL cells harboring the BcTir or BcTpr expression plasmids were cultured in LB medium at 37°C until the optical density at 600 nm (OD _600_) reached 0.8 (for BcTir) or 1.2 (for BcTpr). Protein expression was then induced by 0.1 mM isopropyl β-D-1-thiogalactopyranoside. The cells continued to grow at 16°C (for BcTir) or 12°C (for BcTpr) overnight before harvest. For BcTir, the harvested cells were disrupted in a high-salt lysis buffer (50 mM Tris-HCl (pH 8.0), 25 mM imidazole, 1 M NaCl, 1 mM PMSF and 10 µg/mL DNase I) and subjected to centrifugation. Subsequently, the supernatant was loaded onto a Ni-NTA column. After washes with the lysis buffer, the BcTir protein was eluted with a low-salt buffer (25 mM Tris-HCl (pH 7.5), 300 mM imidazole, 100 mM NaCl), followed by ion-exchange chromatography on a HiTrap Heparin column (GE Healthcare). The BcTir protein was further purified using size-exclusion chromatography on a HiLoad 16/600 Superdex 75 pg column (GE Healthcare) using a buffer containing 20 mM Tris-HCl (pH 7.5), 100 mM NaCl, 5% glycerol, and 5 mM DTT. Our initial purification of BcTpr following a similar approach as described above led to strong protein aggregation as indicated by size-exclusion chromatography. To alleviate this issue, the harvested cells were disrupted in a denaturation buffer containing 50 mM Tris-HCl (pH 8.0), 10 mM imidazole, 1 M NaCl, 6 M guanidine hydrochloride, 10% glycerol, 1 mM PMSF, and 10 µg mL^-1^ DNase I. After centrifugation, the supernatant was loaded onto a Ni-NTA column and the protein was eluted with buffer containing 50 mM Tris-HCl (pH 8.0), 300 mM imidazole, 1 M NaCl, 10% glycerol, and 6 M guanidine hydrochloride. The protein sample was then concentrated and subjected to refolding by dropwise addition into the buffer containing 50 mM Tris-HCl (pH 8.0), 5 mM imidazole, 1 M NaCl, and 5% glycerol. Subsequently, the protein sample was loaded onto another Ni-NTA column, followed by elution using a buffer containing 50 mM Tris-HCl (pH 8.0), 300 mM imidazole, 1 M NaCl, and 5% glycerol. The BcTpr protein was further purified using size-exclusion chromatography on a HiLoad 16/600 Superdex 200 pg column (GE Healthcare) equilibrated with a buffer containing 20 mM Tris-HCl (pH 8.0), 250 mM NaCl, 5% glycerol and 5 mM DTT. Protein purity of BcTir and BcTpr was confirmed using SDS-PAGE. The final protein samples were stored at −80°C for further use.

#### Crystallization, X-ray Data Collection, and Structure Determination

About 1.3 mM of BcTir sample was subjected to sparse-matrix screening to identify initial crystallization conditions (Hampton Research Inc.). The crystals were then reproduced and optimized using the hanging-drop vapor diffusion method with 0.03 M citric acid, 0.07 M BisTris (pH 7.6), and 20% PEG 3,350 at 4°C. The crystals were soaked with cryo-protectant made of the crystallization solution supplemented with 25% glycerol solution before being flash frozen in liquid nitrogen. The X-ray diffraction data of BcTir were collected on the 24-ID-E beamline NE-CAT beamline at the Advanced Photon Source, Argonne National Laboratory. The diffraction data were indexed, integrated and scaled using HKL3000 program (Otwinowski and Minor, 1997). The structure of BcTir was solved by molecular replacement using the PHASER program (McCoy et al., 2007), with the structural model generated using the AlphaFold program as a search model. The initial structural model of BcTir was then subjected to modification using COOT (Emsley and Cowtan, 2004) and refinement using the PHENIX software package (Adams et al., 2002) in an iterative manner. The same R-free test set was used throughout the refinement. The statistics for data collection and structural refinement of the BcTir protein are summarized in Table S4.

#### Measurements of NAD^+^ levels *in vivo*

NAD^+^ levels were assayed as described previously with some modifications (Ofir et al., 2021). Overnight cultures of *E. coli* expressing the BcTir/Tpr system, BcTir, BcTpr, BcTprN, BcTprC, and negative controls were diluted 1:100 into LB supplemented with 50 μg mL^-1^ kanamycin and 50 μM IPTG and grown with shaking at 37°C until an OD _600_ of 0.4. After removing 15 mL of culture for an uninfected control, phage T2 was added at an MOI of 2. Samples were removed after 15, 30, 45 and 60 minutes. The bacteria were centrifuged at 8,000 g for 5 min to pellet. We discarded the supernatant and resuspended the pellet in 300 uL of 1 × phosphate buffer (PBS) supplemented with 4 mg mL^-1^ lysozyme (Sigma, L6876). The sample was transferred to a FastPrep Lysing Matrix B in a 2 mL tube (MP Biomedicals, 116911100) and lysed using a FastPrep bead beater for 20 s at 6 m s^-1^ for two cycles. Tubes were centrifuged at 4°C for 30 min at 16,000 × g. The supernatant was mixed with cold methanol and stored at -20°C until HPLC analysis.

#### Liquid chromatography-mass spectrometry (LC-MS)

The supernatant was mixed with methanol to a final concentration of 75% methanol, and 40 ng of ATP-13^C^10 in 4 µL of 75% methanol was added to 20 µL of the mixture as an internal standard. 10 µL were injected onto the UPLC-MS system for quantification.

Nucleotides were resolved using a HILIC chromatography on a SeQuant ZIC-pHILIC column (5 µM, 150 × 2.1 mm, p/n 504600.001, Sigma-Aldrich, St. Louis, MO) with a ZIC-pHILIC guard pre-column (5 µM, 20 × 2.1 mm, p/n 50438.001, Sigma-Aldrich, St. Louis, MO) at ambient temperature. The LC system consisted of a Waters ACUITY UPLC pump and a well plate autosampler (Waters, Milford, MA). Solvent A was 10 mM ammonium bicarbonate with 0.2% ammonium hydroxide in water, and solvent B was 100% acetonitrile. The flow rate was maintained at 0.2 mL min^-1^. For nucleotide elution, solvent A was increased from 10% to 60% over 8 min, then at 13 min to 90% over 7 min. At 23 min, %A was returned to 10% over 1 min. The equilibration time between injections was 6 min.

MS data acquisition was performed on a quadrupole time-of-flight accurate mass spectrometer (Synapt XS Q-TOF, Waters, Milford, MA) operated in a negative ESI mode. The cone voltage was 40 V and the capillary voltage was 2.0 kV. The source and desolvation temperatures were 120°C and 250°C, respectively. The analyzer was operated with an extended dynamic range at 10,000 resolution (fwhm at m/z 554) with an acquisition time of 0.1 s. MSE mode was used to collect data with the T-wave element alternated between a low energy of 2V and high energy states where the transfer T-wave element voltage was from 10-25 V(Brose et al., 2013). The cone gas flow rate was 10 L h^-1^ and the desolvation gas flow was 500 L h^-1^. Leucine enkephalin (200 pg μL^-1^, ACN: water, 50: 50 by volume) was infused at a rate of 10 μL min^-1^ for mass correction.

For quantification, extracted ion chromatograms for ATP-13^C^10 (m/z 516.022), NAD^+^ (662.101), and cADPR (m/z 540.053) using a 10 mDa mass window and integrated using QuanLynx. Peak identification was performed based on exact mass, retention time, and high energy (MS/MS) spectra compared to standards. Nucleotides were quantified using a standard curve built against ATP-13^C^10 internal standard. MassLynx V4.2 software (Waters) was used for instrument control, acquisition, and sample analysis.

#### NADase assay *in vitro*

NADase assays were performed as previously described (Ka et al., 2020). Purified BcTir, BcTpr, BcTprN, and BcTprC proteins (2.5 μM) and NAD^+^ (250 μM) were incubated in reaction buffer (63 mM NaCl, 14 mM Tris-HCl, pH 7.5) in a 100 μL final reaction volume. Reactions were carried out at 30°C for the indicated amount of time. Reactions were stopped by the addition of cold methanol and stored at -20°C for at least 1 h to precipitate. The samples were centrifuged at 4°C for 30 min at 16,000 × g. After centrifugation of the samples, 20 μL of the supernatants were mixed with 4 μL ATP-^13^C^10^ for LC-MS analysis.

#### Flow cytometry analysis of infected cells

*E.coli* BL21 (DE3) strains carrying recombinant plasmids BcTir/Tpr and empty vector pET28a were grown overnight at 37°C and diluted in LB medium supplemented with 50 μg mL^-1^ kanamycin and 50 μM IPTG at 37°C to an OD _600_ of 0.4. The cultures were infected with phage T2 at an MOI of 2. After 45 min infection, each culture was diluted into filtered PBS containing 1 μL of propidium iodide. The diluted bacteria were incubated in the dark at room temperature for 5 minutes and filtered by using a 40-mm filter. The cells were analyzed by a BD FACSymphony^TM^ A3 (San Jose, CA, USA) and analyzed by FlowJo v10.8.1 software.

#### Transmission electron microscopy of bacterial cells

The samples were collected as mentioned above. After removing 200 μL of culture for an uninfected control, phage T2 was added at an MOI of 2. Samples were removed after 15, 30, 45, and 60 minutes. Samples were centrifuged at 8,000 × g for 2 min. The pellet was fixed with 2% paraformaldehyde/2% glutaraldehyde in cacodylate buffer at RT for 1 h, followed by 3 rinses with 2 mL of 0.1 M sodium cacodylate buffer (pH 7.4). Samples were post-fixed with 1% OsO_4_/1.5% potassium ferrocyanide in 0.1 M cacodylate buffer (pH 7.4) on ice for 1 h then washed 3 times with distilled water. We dehydrated samples with a graded series of ethanol solutions and infiltrated them overnight in a 1:1 mixture of EMBed 812:100% ethanol. We then transferred samples to 100% EMBed, incubated them all day, removed EMBed, replaced with fresh EMBed, and cured them for 48 h at 60°C. We finally cut the bottoms from the dishes, separated the embedding resin from the plastic bottom of the dish by plunging into liquid nitrogen and reembedded in sample molds for sectioning. Samples were sectioned using an RHC MTX Ultramicrotome and a diatom diamond knife and then placed on 8 GC 180 copper grids. These grids were viewed on a Hitachi 7500H Transmission Electron Microscope. Images were taken and analyzed according to our previously published methods (Wu et al., 2005).

#### Adsorption and DNA replication assay

The samples were collected as mentioned above. T2 phage were added at an MOI of 2. An aliquot of 500 μL of culture was taken at 0, 15, 20, 30, 45, and 60 min after infection. The bacteria were centrifuged at 8,000 g for 3 min to pellet. Supernatants were used for the adsorption assay and the pellet was used for the DNA replication assay. Total DNA was extracted from the supernatant and pelleted using the chloroform–phenol method. Absolute quantification PCR was used to quantify the copy number of phage DNA. The T2 phage-specific sequence was amplified by using primers (YLCZ047 and YLCZ048) and purified for standard template construction and converted the amount of DNA (ng) into copy of numbers, and followed a dilution course of tenfold, from 10^9^ to 10^1^ per microliter. Primers (YLCZ049 and YLCZ050) specifically targeting the gene sequence of Enterobacteria phage T2 was used. The reactions were performed in triplicate using a Thermal Cycler (CFX96 Real-Time sytem, Bio-Rad Laboratories, Hercules, CA, USA). All the sequences and primers are listed in Supplementary Table 2.

#### Measurements of ROS and antioxidant capability in *E. coli*

The samples were collected as mentioned above. After removing 10 mL culture for an uninfected control, T2 phage was added at an MOI of 2. Samples were removed after 15 and 30 minutes. For ROS, cells were harvested by concentration, washed twice, resuspended in 1 × PBS containing 8 μM of the dye CM-H_2_DCFDA. The cultures were incubated at 37°C for 30 minutes and fluorescence was measured using a FlexStation® 3 Multi-Mode Microplate Reader (Sunnyvale, CA, USA) with a 485 nm excitation and a 528-nm emission filter. The antioxidant activity was measured by an antioxidant capacity kit according to the manufacturer’s instructions. Proteins were extracted using the B-PER reagent. The extracts were clarified by centrifugation (14,000 g at 4°C for 30 min). A working solution was freshly prepared by mixing 100 μL of samples and the Cu^2+^ working solution and incubating the reaction for 90 minutes at room temperature. The plate was protected from light during the incubation. The absorbance of the reaction mixture was recorded at 570 nm. A standard curve was generated using a Trolox standard at concentrations ranging from 4 to 20 nM. The Trolox equivalents of the samples were calculated from the standard curve.

#### RNA isolation and quantitative reverse-transcriptional PCR

Total RNA was isolated using RNeasy Mini Kit according to the manufacturer’s protocol with DNase I treatment. 0.5 μg total DNA-free RNA was reverse transcribed to cDNA using the PrimeScript™ RT reagent Kit with gDNA Erase. qPCR was performed using SsoAdvanced Universal SYBR Green Supermix and run at CFX ConnectTM Real-Time System (Bio-Rad). The 16S rRNA gene was used as the endogenous reference control to normalize for differences in total RNA quantity, and relative gene expression was determined using the comparative threshold cycle 2^−ΔΔCT^ method (Livak and Schmittgen, 2001).

#### Pull-down assay

The BcTir was amplified from the pET28a-BcTir and ligated into the pPGH plasmid (Seeburg *et al*., 1983) between the NdeI and XhoI restriction sites using the in-fusion snap assembly master mix kit. This recombinant plasmid was first transformed into *E. coli* One Shot™ TOP10 and then introduced into *E. coli* host strain BL21 (DE3). Primers are listed in Supporting Table S2. The BcTir-GST was expressed and purified using the Pierce® Glutathione Agarose. The pull-down assay was performed using a Pierce^TM^ Pull-down PolyHis Protein:Protein Interaction Kit. The PcTpr, BcTprN, and BcTprC were used as bait proteins and BcTir-GST was used as the prey protein. Recombinant GST epitope tag His Protein (Novus Biologicals) was used as the negative control. Briefly, the HisPur Cobalt Resin was incubated with 100 μg BcTpr protein for 2 h with gentle rocking at 4°C. After washing five times, 100 μg BcTir-GST protein was added and incubated at 4°C overnight with gentle rocking. We collected the flow through from the resin and washed it five times. Bait-Prey protein was eluted with 290 mM imidazole in the wash solution. The eluted proteins were analyzed by SDS-PAGE.

#### Bio-layer interferometry (BLI) analysis

The BLItz™ Label-Free Protein Analysis System (ForteBio, CA, USA) was used for biolayer interferometry, which is a label-free technique and measures protein:protein interactions. The purified BcTpr protein (30 μg) was used as the immobilized protein and was diluted in the assay buffer (63 mM NaCl, 14 mM Tris-HCl, pH 7.5) in the 200 μL final reaction volume. The BcTpr protein was immobilized on the biosensor tip surface using Octet® Ni-NTA (NTA) Biosensors (Sartorius,18-5101) onto which the BcTir-GST protein (BcTpr:BcTir-GST, molar ratio is 1:1) was loaded. Data analyses were performed using GraphPad Prism 9.3.0 software.

#### ITC measurements

Isothermal Titration Calorimetry (ITC) measurements were obtained using a MicroCal iTC200 instrument (GE Healthcare). To measure the bindings of full-length BcTpr with cADPR (Sigma-Aldrich, C7344) and/or NAD^+^ (Sigma-Aldrich, NAD100-RO), BcTpr was subjected to overnight dialysis against buffer containing 100 mM NaCl, 25 mM Tris-HCl (pH 7.4), and 1 mM 2-mercaptoethanol. 0.05 mM BcTpr was first titrated with 0.5 mM cADPR, then with 0.5 mM NAD^+^ at 20°C. Before the titration, buffer-to-buffer measurements were performed to ensure low background error of baseline. Origin 7.0 software was used for data analysis. A single-site binding mode was used for fitting.

#### Proteins alignment and phylogenetic analysis

Protein sequences were aligned by Clustal omega (Sievers et al., 2011) and the output was visualized by ESPript 3.0 (Robert and Gouet, 2014) combined with the second structures. There were 78 Tpr prokaryotic homologs that were retrieved from the previous database (Burroughs and Aravind, 2020). The phylogenetic tree was constructed based on the alignment of Tpr proteins via Clustal Omega. The output tree file was visualized and modified by ggtree (Yu et al., 2017).

#### Data analyses

Graph plotting and statistical analyses were performed using GraphPad Prism 9.3.0 (Graph-Pad Software). Comparisons were analyzed by unpaired parametric t-test, two-tailed with no corrections.

## Supplemental figures

**Figure S1.**
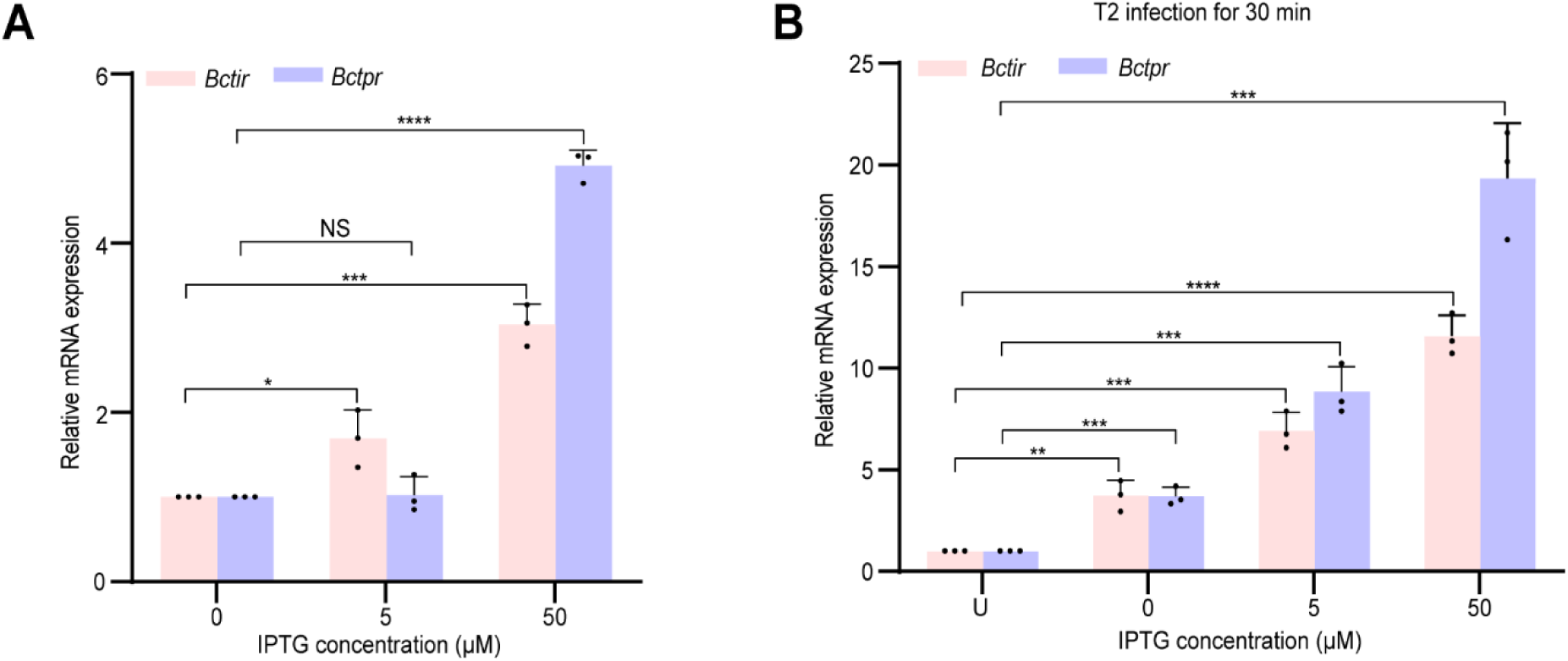
The expression of the BcTir/Tpr system induced by different IPTG concentrations under normal conditions (A) and phage infection (B). The relative expression of the genes, *Bctir* and *Bctpr*, were measured by reverse transcription quantitative PCR following a concentration course of IPTG at 0 µM, 5 µM and 50 µM without (A) and with phage infection (B). “U” in B indicates uninfection. Three independent replicates were used for calculating the mean values and error bars represent the standard error of the mean, SEM. Data from A and B were analyzed compared to the “without IPTG” (A) and “uninfection” (B) by using two-tailed student’s *t*-test. “****” indicates *p-*value < 0.0001, “***” indicates *p-*value < 0.001, “**” indicates *p-*value < 0.01, and “*” indicates *p*-value < 0.05. “NS” means no significance.

**Figure S2.**
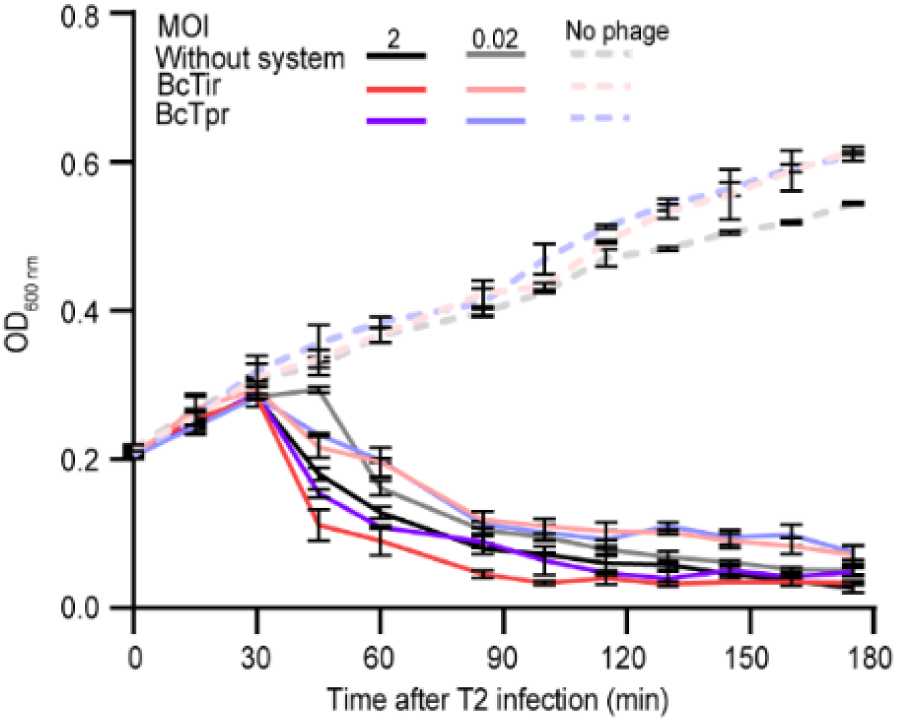
Cell growth of the *E.coli* carrying the BcTir, BcTpr and none at MOI dilutions of 2, 0.02, and 0. Three independent replicates were performed for each sample. Data are presented as mean ± SEM.

**Figure S3.**
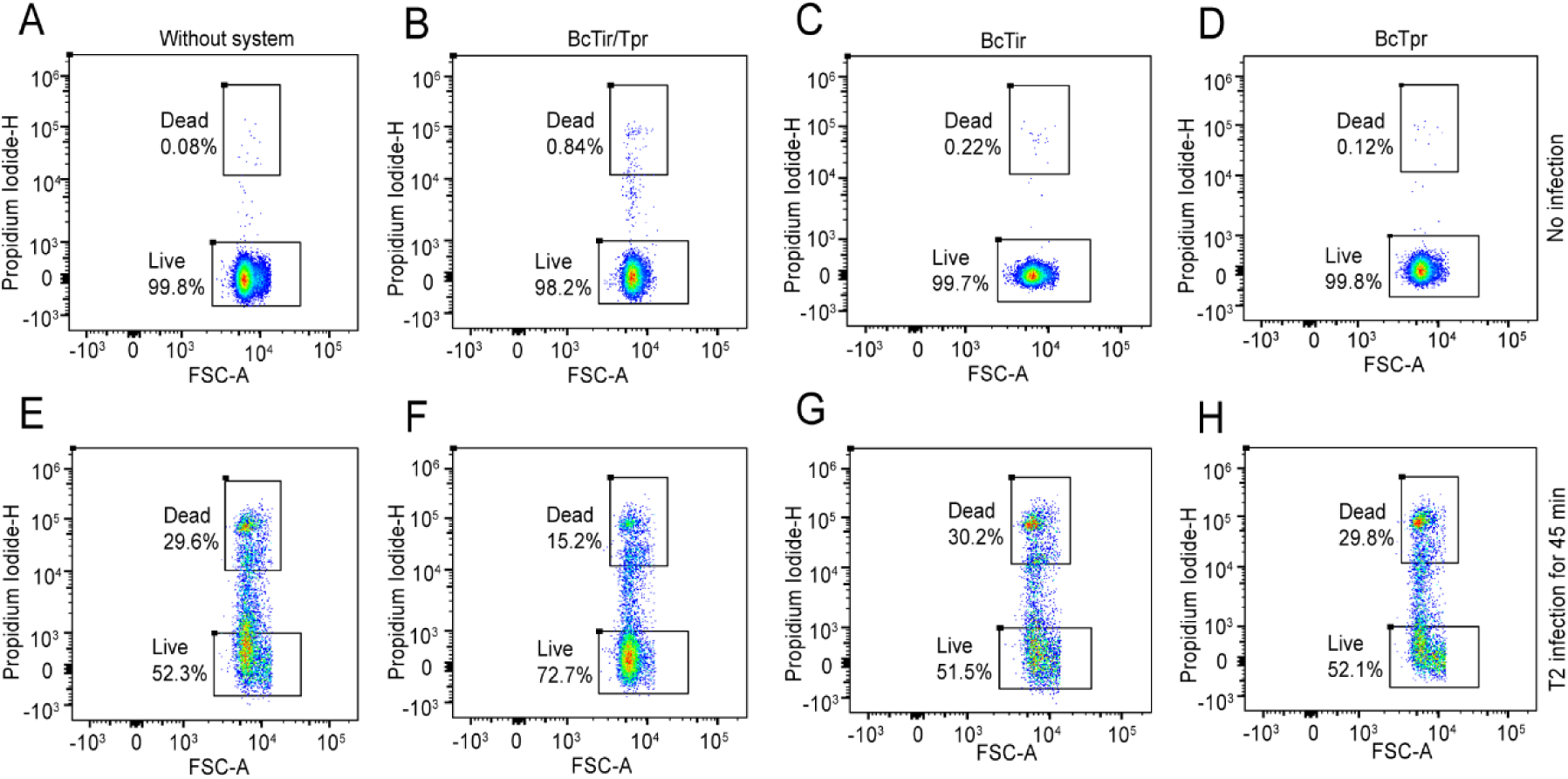
Fluorescence activated cell sorting (FACS) was used to distinguish living and dead cells. Plots were gated on forward scatter (FSC-A) and side scatter (Propidium Iodide-H) to define cell survival of transformants expressing the empty vector, BcTir/Tpr system, BcTir, BCTpr without (A, B, C and D) and with (E, F, G, and H) phage infection compared to control cells at 45 min post infection.

**Figure S4.**
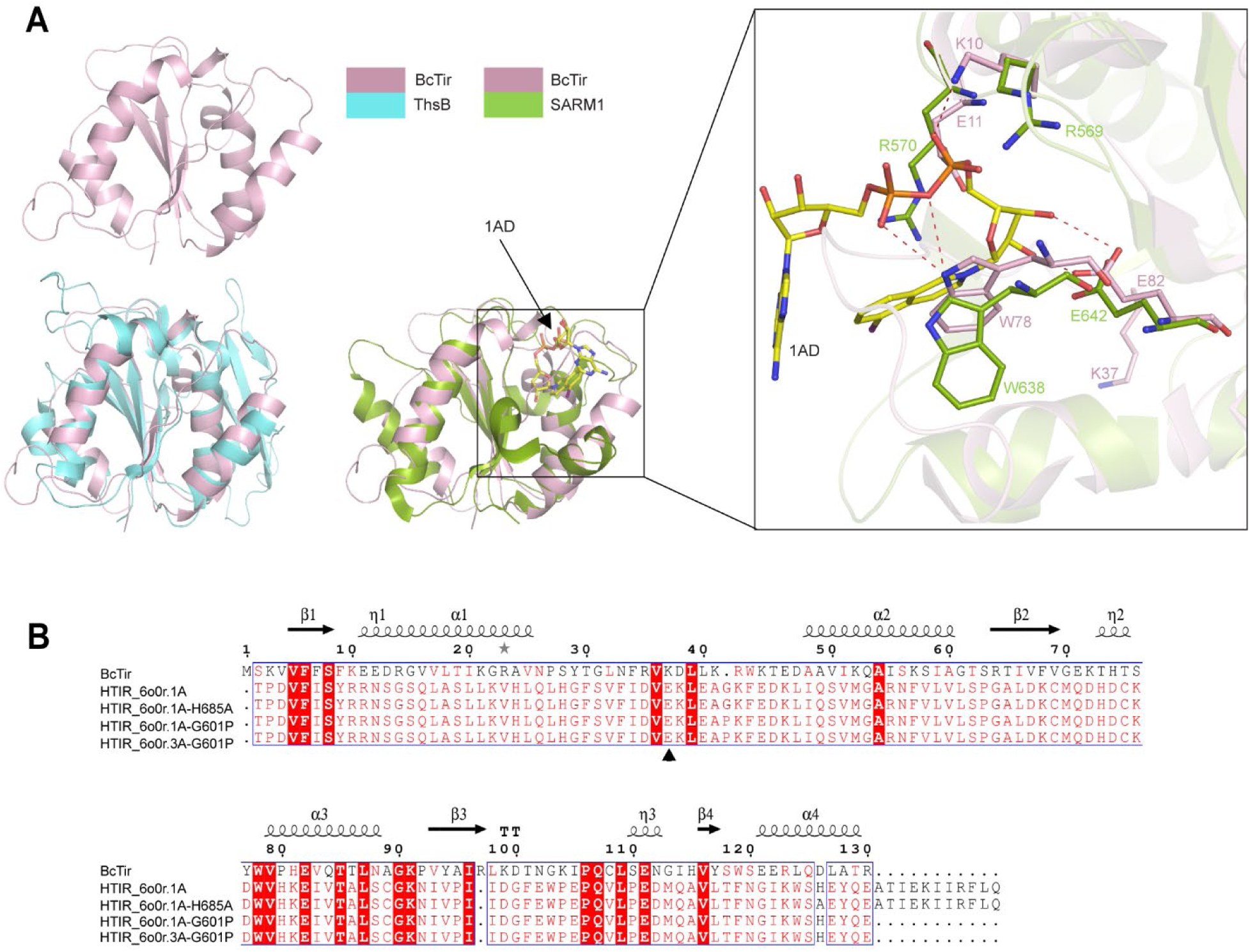
Structural superimposition and sequence alignment of BcTir. (A) Crystal structure of BcTir and its superposition with those of ThsB in the thoeris system (PDB: 6HLY) and HTIR_SARM1 (PDB: 7NAK) bound to the inhibitor iodine isoquinoline adenine dinucleotide (1AD). Potential residues of BcTir, K10, E11, and E82, were assumed to form hydrogen bonds with 1AD. (B) Structure-based sequence alignment between the BcTir and the TIR domain variants of human SARM1 (hSARM1). Arrow indicates the catalytic site, glutamate (E) of the TIR domain of human SARM1, corresponding to residue lysine (K) at 37 of BcTir.

**Figure S5.**
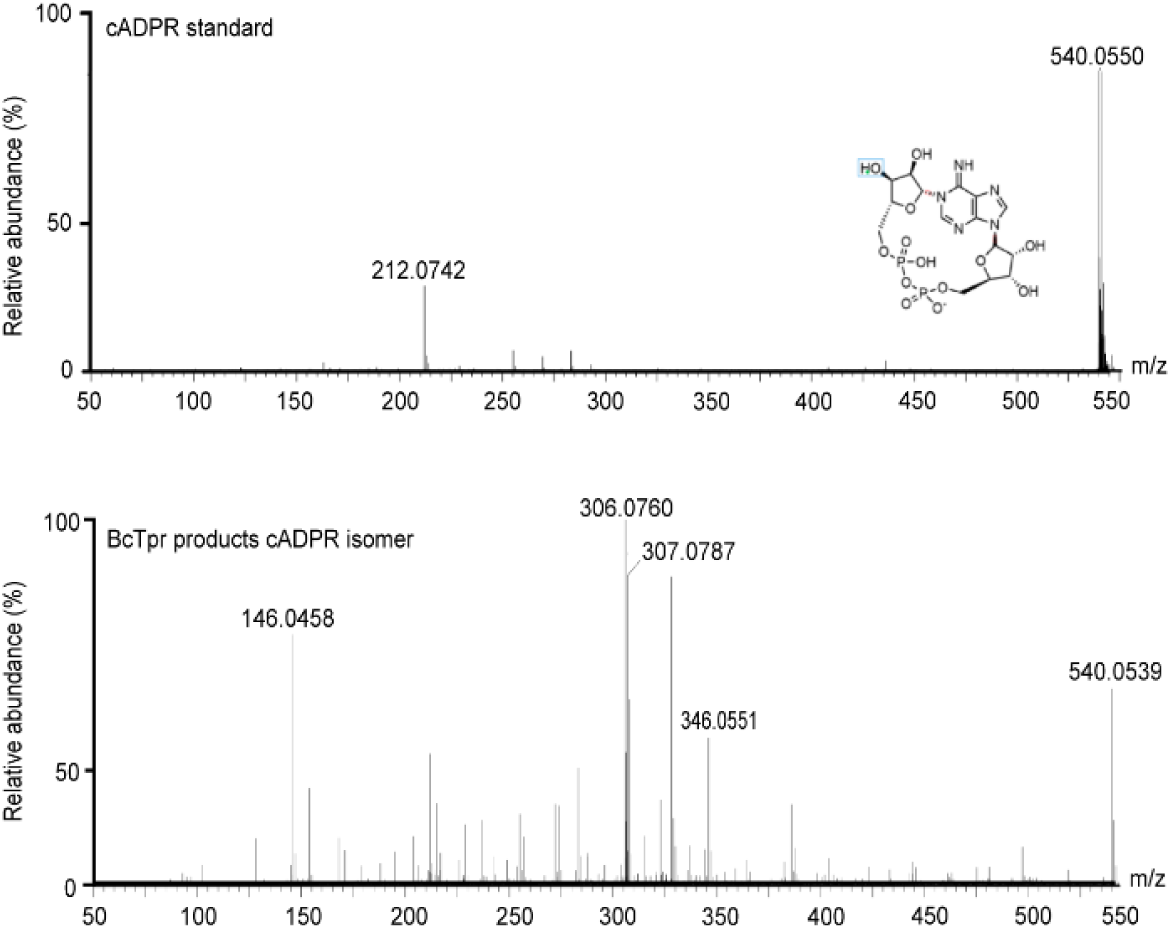
MS/MS fragmentation spectra of standard cADPR (Top) and BcTpr-generated cADPR isomer (bottom). Hypothesized structures of MS/MS fragments of cADPR are presented.

**Figure S6.**
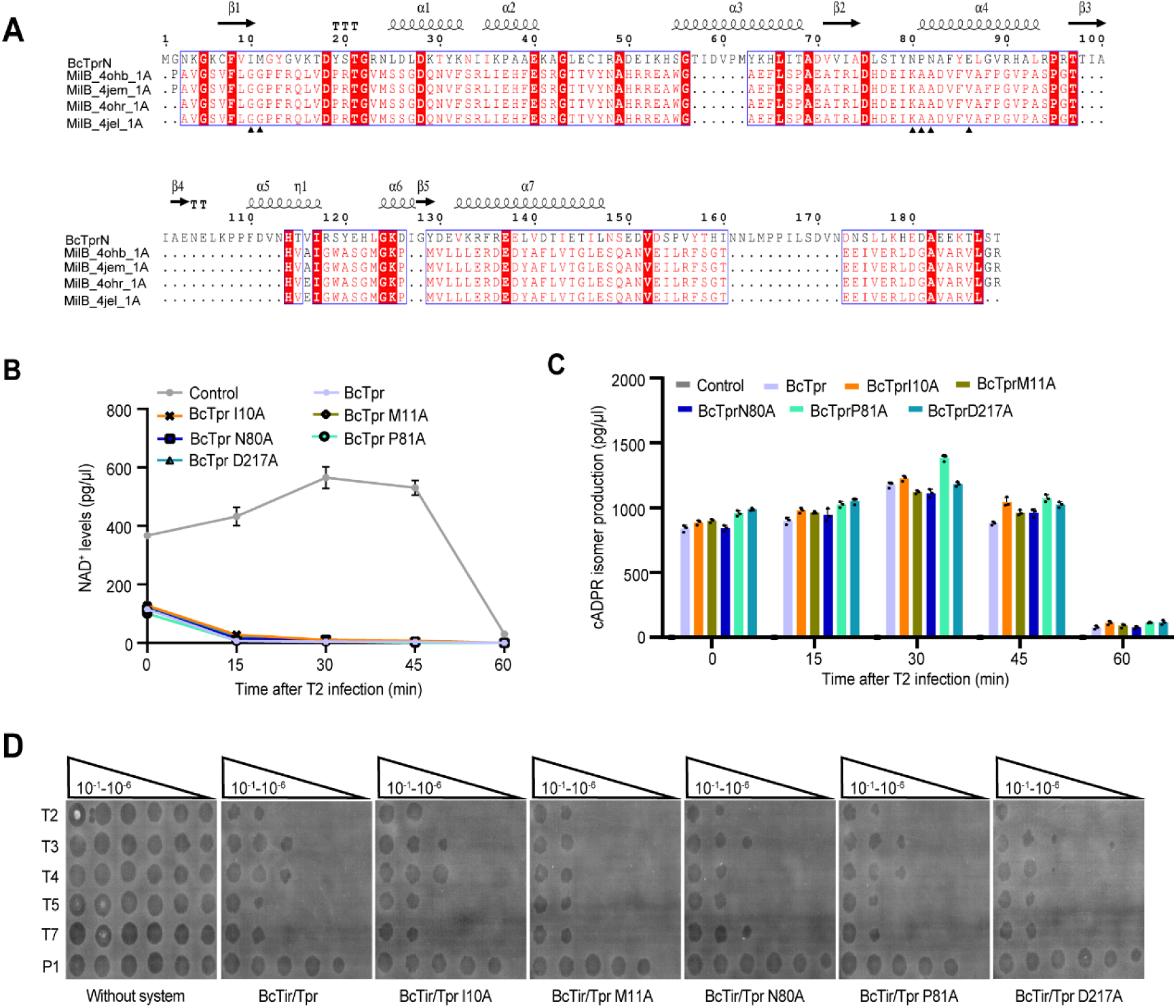
Potential residues tested for anti-phage immunity and NAD^+^ metabolism. (A) Structure-based sequence alignment of BcTprN to MilB in various structural configurations. Arrows below the alignment indicate the potential NAD^+^ binding residues of BcTprN, excluding D217, from BcTprC. (B) Site mutations in BcTpr I10A, BcTpr M11A, BcTpr N80A, BcTpr P81A and BcTpr D217A, were not the critical catalytical sites for NAD^+^ consumption. There was also no cADPR isomer production following these site mutations shown in C. (D) These site mutations were unable to abolish the anti-phage immunity conferred by BcTir/Tpr system. The data from B and C were obtained from three independent replicates and the data are presented as mean ± SEM.

**Figure S7.**
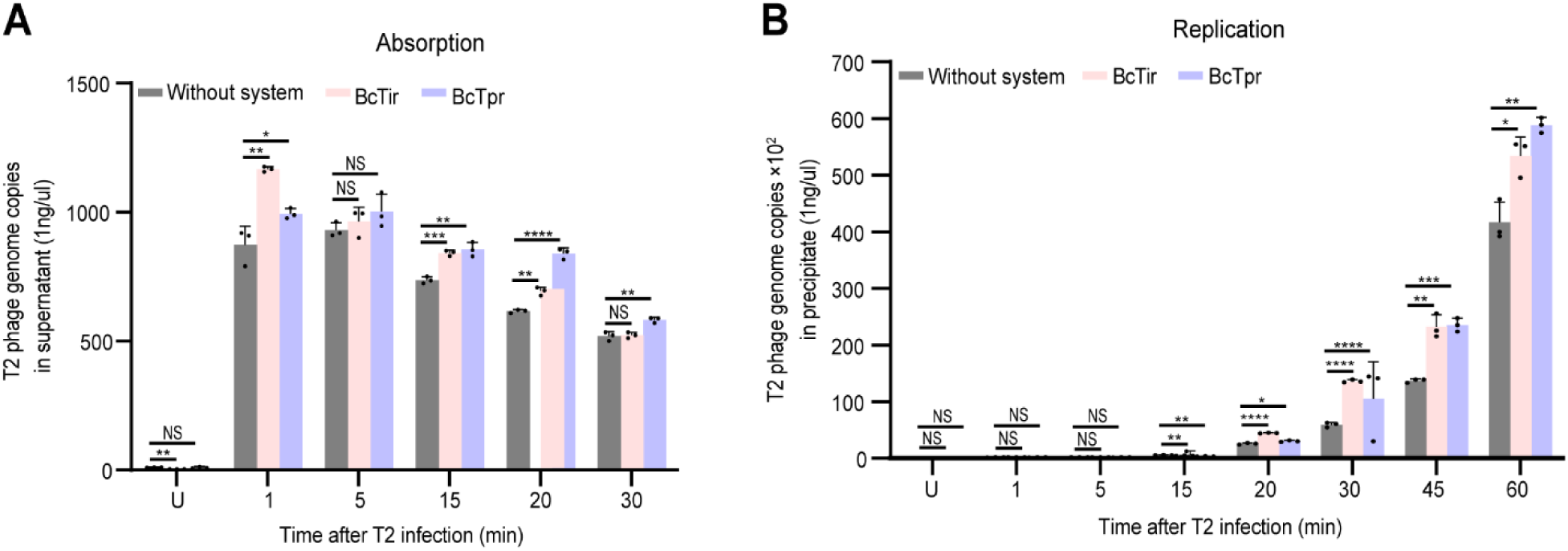
The influences of BcTir and BcTpr on phage adsorption and replication. (A) Phage adsorption was assessed by detecting the copy numbers of phage in the supernatant via qPCR during the time course of 30 min post phage infection. More phage copy numbers in the supernatant indicated inability of phage entry or adsorption. (B) By contrast, the precipitated population was calculated to represent phage replication during the time course after phage infection. “U” in A, B indicates uninfection. The data were obtained from three independent replicates and the data are presented as mean ± SEM. Data from A and B were analyzed compared to the “Without system” at the same time point by using two-tailed student’s *t*-test. “****” indicates *p-*value < 0.0001, “***” indicates *p-*value < 0.001, “**” indicates *p-*value < 0.01, and “*” indicates *p*-value < 0.05. “NS” means no significance.

**Figure S8.**
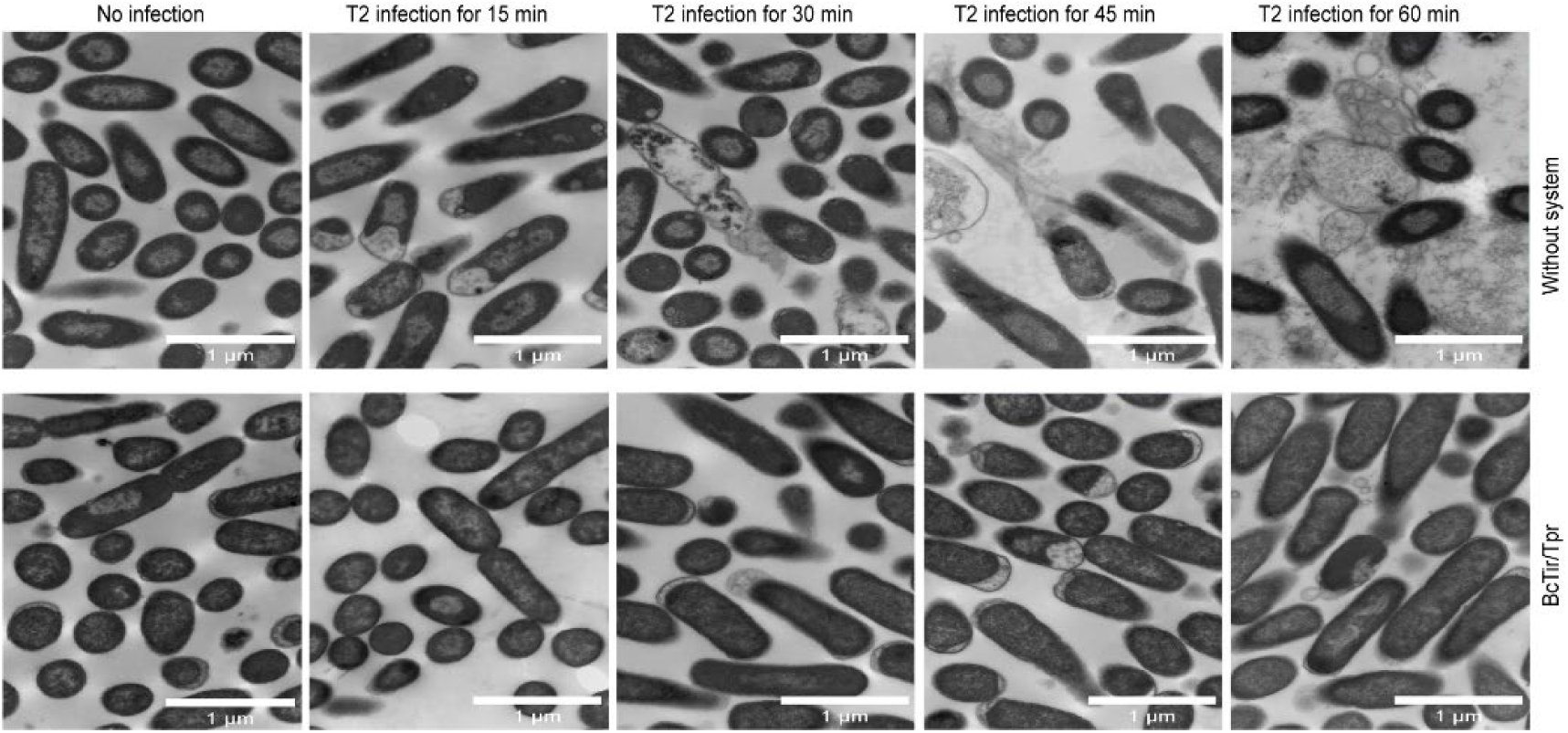
Transmission electronic for detecting membrane dynamics following a time course of phage infection. Observations from TEM of a 60-minute time course for *E. coli* with and without the BcTir/Tpr system after infection with phage T2 at the MOI of 2.

**Figure S9.**
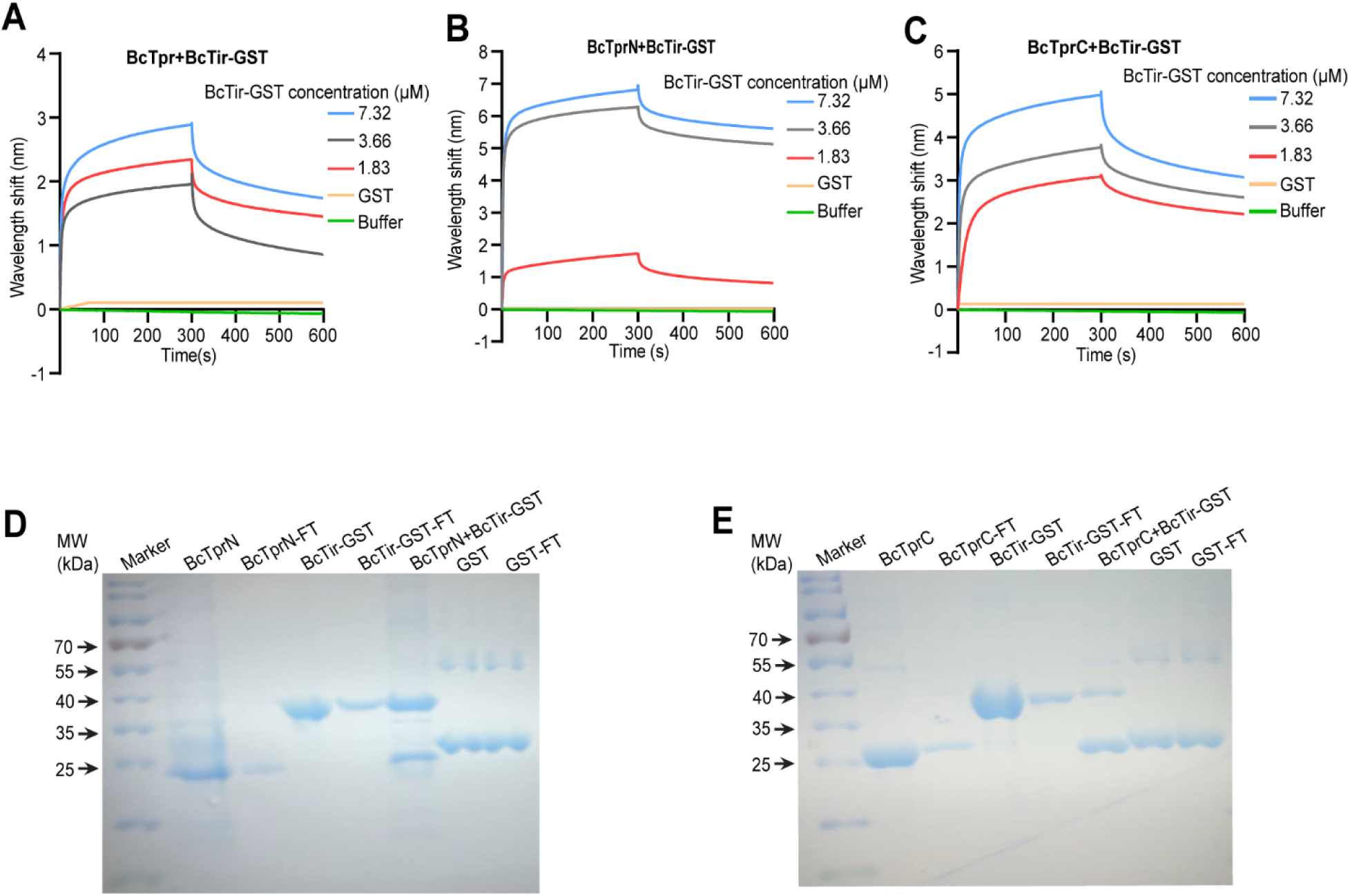
BcTir interacted with BcTpr at both the N terminus and C terminus detected by Bio-layer interferometer (BLI) and pull-down assays. (A) BcTir interacts with BcTpr detected by BLI in a concentration course. (B) and (D), BcTir interacts with BcTprN detected by BLI and pull-down assays. (C) and (E), BcTir interacts with BcTprC measured by BLI and pull-down assays. GST as the tag for BcTir, was checked as the negative control to rule out its non-specific interaction with BcTpr. Marker: protein ladder of different molecular weights; FT: flow through.

**Figure S10.**
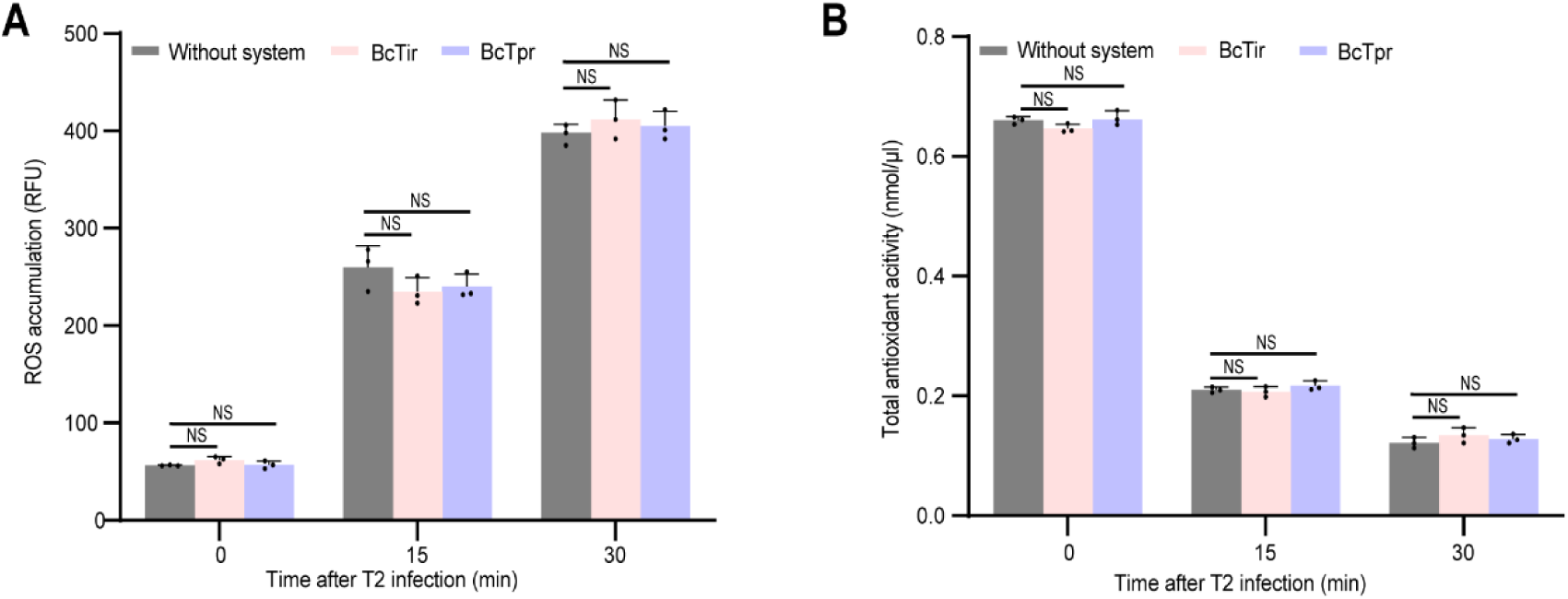
The BcTir and BcTpr can’t protect cell viability from ROS oxidation induced by phage infection. Cellular ROS accumulation (A) and total antioxidant activity (B) of cells with or without BcTir and BcTpr were measured at 0, 15, and 30 min after T2 phage infection. The data were obtained from three independent replicates and the data are presented as mean ± SEM. Data from A and B were analyzed compared to the “Without system” at the same time point by using two-tailed student’s *t*-test. “NS” means no significance.

## References

Abbas, Y.M., Laudenbach, B.T., Martinez-Montero, S., Cencic, R., Habjan, M., Pichlmair, A., Damha, M.J., Pelletier, J., and Nagar, B. (2017). Structure of human IFIT1 with capped RNA reveals adaptable mRNA binding and mechanisms for sensing N1 and N2 ribose 2’-O methylations. Proc. Natl. Acad. Sci. USA. 114, E2106–E2115.

Abbas, Y.M., Pichlmair, A., Gorna, M.W., Superti-Furga, G., and Nagar, B. (2013). Structural basis for viral 5’-PPP-RNA recognition by human IFIT proteins. Nature 494, 60–64.

Adams, P.D., Grosse-Kunstleve, R.W., Hung, L.-W., Ioerger, T.R., McCoy, A.J., Moriarty, N.W., Read, R.J., Sacchettini, J.C., Sauter, N.K., and Terwilliger, T.C. (2002). PHENIX: building new software for automated crystallographic structure determination. Acta Crystallogr. D Biol. Crystallogr. 58, 1948–1954.

Allan, R.K., and Ratajczak, T. (2011). Versatile TPR domains accommodate different modes of target protein recognition and function. Cell Stress Chaperones. 16, 353–367.

Boatright, K.M., and Salvesen, G.S. (2003). Mechanisms of caspase activation. Curr. Opin. Cell. Biol. 15, 725–731.

Brose, S.A., Baker, A.G., and Golovko, M.Y. (2013). A fast one-step extraction and UPLC– MS/MS analysis for E_2_/D_2_ Series prostaglandins and isoprostanes. Lipids 48, 411–419.

Brzozowski, R.S., Huber, M., Burroughs, A.M., Graham, G., Walker, M., Alva, S.S., Aravind, L., and Eswara, P.J. (2019). Deciphering the role of a SLOG superfamily protein YpsA in Gram-positive bacteria. Front. Microbiol. 10, 623.

Burroughs, A.M., and Aravind, L. (2020). Identification of uncharacterized components of prokaryotic immune systems and their diverse eukaryotic reformulations. J. Bacteriol 202, e00365–00320.

Burroughs, A.M., Zhang, D., Schäffer, D.E., Iyer, L.M., and Aravind, L. (2015). Comparative genomic analyses reveal a vast, novel network of nucleotide-centric systems in biological conflicts, immunity and signaling. Nucleic Acids Res. 43, 10633–10654.

Coronas-Serna, J.M., Louche, A., Rodríguez-Escudero, M., Roussin, M., Imbert, P.R., Rodríguez-Escudero, I., Terradot, L., Molina, M., Gorvel, J.-P., and Cid, V.J. (2020). The TIR-domain containing effectors BtpA and BtpB from *Brucella abortus* impact NAD metabolism. PLOS Pathog. 16, e1007979.

Cui, N., Zhang, J.-T., Li, Z., Liu, X.-Y., Wang, C., Huang, H., and Jia, N. (2022). Structural basis for the non-self RNA-activated protease activity of the type III-E CRISPR nuclease-protease Craspase. Nat. Commun 13, 1–13.

D’Andrea, L.D., and Regan, L. (2003). TPR proteins: the versatile helix. Trends. Biochem. Sci. 28, 655–662.

D’Autréaux, B., and Toledano, M.B. (2007). ROS as signalling molecules: mechanisms that generate specificity in ROS homeostasis. Nat. Rev. Mol. Cell Biol. 8, 813–824. 10.1038/nrm2256.

Dong, T.G., Dong, S., Catalano, C., Moore, R., Liang, X., and Mekalanos, J.J. (2015). Generation of reactive oxygen species by lethal attacks from competing microbes. Proc. Natl. Acad. Sci. USA 112, 2181–2186.

Dvorak, P., Chrast, L., Nikel, P.I., Fedr, R., Soucek, K., Sedlackova, M., Chaloupkova, R., de Lorenzo, V., Prokop, Z., and Damborsky, J. (2015). Exacerbation of substrate toxicity by IPTG in *Escherichia coli* BL21 (DE3) carrying a synthetic metabolic pathway. Microb. Cell Fact. 14, 1–15.

Eastman, S., Smith, T., Zaydman, M.A., Kim, P., Martinez, S., Damaraju, N., DiAntonio, A., Milbrandt, J., Clemente, T.E., and Alfano, J.R. (2022). A phytobacterial TIR domain effector manipulates NAD^+^ to promote virulence. New Phytol. 233, 890–904.

Einsfeldt, K., Júnior, J.B.S., Argondizzo, A.P.C., Medeiros, M.A., Alves, T.L.M., Almeida, R.V., and Larentis, A.L. (2011). Cloning and expression of protease ClpP from *Streptococcus pneumoniae* in *Escherichia coli*: study of the influence of kanamycin and IPTG concentration on cell growth, recombinant protein production and plasmid stability. Vaccine 29, 7136–7143.

Emsley, P., and Cowtan, K. (2004). Coot: model-building tools for molecular graphics. Acta Crystallogr. D Biol. Crystallogr. 60, 2126–2132.

Essuman, K., Milbrandt, J., Dangl, J.L., and Nishimura, M.T. (2022). Shared TIR enzymatic functions regulate cell death and immunity across the tree of life. Science 0, eabo0001.

Essuman, K., Summers, D.W., Sasaki, Y., Mao, X., DiAntonio, A., and Milbrandt, J. (2017). The SARM1 toll/interleukin-1 receptor domain possesses intrinsic NAD^+^ cleavage activity that promotes pathological axonal degeneration. Neuron 93, 1334–1343. e1335.

Essuman, K., Summers, D.W., Sasaki, Y., Mao, X., Yim, A.K.Y., DiAntonio, A., and Milbrandt, J. (2018). TIR domain proteins are an ancient family of NAD^+^-consuming enzymes. Curr. Biol. 28, 421–430 e424.

Huang, S., Jia, A., Song, W., Hessler, G., Meng, Y., Sun, Y., Xu, L., Laessle, H., Jirschitzka, J., Ma, S., et al. (2022). Identification and receptor mechanism of TIR-catalyzed small molecules in plant immunity. Science 0, eabq3297.

Huiting, E., and Bondy-Denomy, J. (2023). Defining the expanding mechanisms of phage-mediated activation of bacterial immunity. Curr Opin Microbiol. 74, 102325.

Jia, A., Huang, S., Song, W., Wang, J., Meng, Y., Sun, Y., Xu, L., Laessle, H., Jirschitzka, J., Hou, J., et al. (2022). TIR-catalyzed ADP-ribosylation reactions produce signaling molecules for plant immunity. Science 0, eabq8180.

Johnson, A.G., Wein, T., Mayer, M.L., Duncan-Lowey, B., Yirmiya, E., Oppenheimer-Shaanan, Y., Amitai, G., Sorek, R., and Kranzusch, P.J. (2022). Bacterial gasdermins reveal an ancient mechanism of cell death. Science 375, 221–225.

Jumper, J., Evans, R., Pritzel, A., Green, T., Figurnov, M., Ronneberger, O., Tunyasuvunakool, K., Bates, R., Žídek, A., and Potapenko, A. (2021). Highly accurate protein structure prediction with AlphaFold. Nature 596, 583–589.

Ka, D., Oh, H., Park, E., Kim, J.H., and Bae, E. (2020). Structural and functional evidence of bacterial antiphage protection by Thoeris defense system via NAD^+^ degradation. Nat. Commun 11, 2816.

Koopal, B., Potocnik, A., Mutte, S.K., Aparicio-Maldonado, C., Lindhoud, S., Vervoort, J.J., Brouns, S.J., and Swarts, D.C. (2022). Short prokaryotic Argonaute systems trigger cell death upon detection of invading DNA. Cell 185, 1471–1486.

Liu, Y., Zhang, C., Wang, Z., Lin, M., Wang, J., and Wu, M. (2021). Pleiotropic roles of late embryogenesis abundant proteins of *Deinococcus radiodurans* against oxidation and desiccation. Comput. Struct. Biotechnol. J. 19, 3407–3415.

Liu, Y.W., Han, C.H., Lee, M.H., Hsu, F.L., and Hou, W.C. (2003). Patatin, the tuber storage protein of potato (*Solanum tuberosum* L.), exhibits antioxidant activity in vitro. J. Agric. Food. Chem. 51, 4389–4393.

Livak, K.J., and Schmittgen, T.D. (2001). Analysis of relative gene expression data using real-time quantitative PCR and the 2^−ΔΔCT^ method. Methods 25, 402–408.

Lopatina, A., Tal, N., and Sorek, R. (2020). Abortive infection: bacterial suicide as an antiviral immune strategy. Annu. Rev. Virol. 7, 371–384.

Mazzocco, A., Waddell, T.E., Lingohr, E., and Johnson, R.P. (2009). Enumeration of bacteriophages using the small drop plaque assay system. In Bacteriophages: methods and protocols, (Springer), pp. 81–85.

McCoy, A.J., Grosse-Kunstleve, R.W., Adams, P.D., Winn, M.D., Storoni, L.C., and Read, R.J. (2007). Phaser crystallographic software. J Appl Crystallogr 40, 658–674.

Ofir, G., Herbst, E., Baroz, M., Cohen, D., Millman, A., Doron, S., Tal, N., Malheiro, D.B.A., Malitsky, S., Amitai, G., and Sorek, R. (2021). Antiviral activity of bacterial TIR domains via immune signalling molecules. Nature 600, 116–120.

Otwinowski, Z., and Minor, W. (1997). Processing of X-ray diffraction data collected in oscillation mode. Method Enzymol 276, 307–326.

Prodromou, C., Siligardi, G., O’Brien, R., Woolfson, D.N., Regan, L., Panaretou, B., Ladbury, J.E., Piper, P.W., and Pearl, L.H. (1999). Regulation of Hsp90 ATPase activity by tetratricopeptide repeat (TPR)-domain co-chaperones. EMBO J. 18, 754–762.

Ramirez, D.H., Yang, B., D’Souza, A.K., Shen, D., and Woo, C.M. (2021). Truncation of the TPR domain of OGT alters substrate and glycosite selection. Anal Bioanal Chem 413, 7385–7399.

Robert, X., and Gouet, P. (2014). Deciphering key features in protein structures with the new ENDscript server. Nucleic Acids Res 42, W320–W324.

Rydel, T.J., Williams, J.M., Krieger, E., Moshiri, F., Stallings, W.C., Brown, S.M., Pershing, J.C., Purcell, J.P., and Alibhai, M.F. (2003). The Crystal Structure, Mutagenesis, and Activity Studies Reveal that Patatin Is a Lipid Acyl Hydrolase with a Ser-Asp Catalytic Dyad. Biochemistry 42, 6696–6708.

Scheufler, C., Brinker, A., Bourenkov, G., Pegoraro, S., Moroder, L., Bartunik, H., Hartl, F.U., and Moarefi, I. (2000). Structure of TPR domain–peptide complexes: critical elements in the assembly of the Hsp70–Hsp90 multichaperone machine. Cell 101, 199–210.

Seeburg, P.H., Sias, S., Adelman, J., de Boer, H.A., Hayflick, J., Jhurani, P., Goeddel, D.V., and Heyneker, H.L. (1983). Efficient bacterial expression of bovine and porcine growth hormones. DNA 2, 37–45.

Seed, K.D. (2015). Battling phages: how bacteria defend against viral attack. PLoS Pathog 11, e1004847.

Shi, Y., Kerry, P.S., Nanson, J.D., Bosanac, T., Sasaki, Y., Krauss, R., Saikot, F.K., Adams, S.E., Mosaiab, T., Masic, V., et al. (2022). Structural basis of SARM1 activation, substrate recognition, and inhibition by small molecules. Mol. Cell 82, 1643–1659.e1610.

Sievers, F., Wilm, A., Dineen, D., Gibson, T.J., Karplus, K., Li, W., Lopez, R., McWilliam, H., Remmert, M., and Söding, J. (2011). Fast, scalable generation of high-quality protein multiple sequence alignments using Clustal Omega. Mol. Syst. Biol. 7, 539.

Sikowitz, M.D., Cooper, L.E., Begley, T.P., Kaminski, P.A., and Ealick, S.E. (2013). Reversal of the substrate specificity of CMP N-glycosidase to dCMP. Biochemistry 52, 4037–4047.

Song, H.K., Sohn, S.H., and Suh, S.W. (1999). Crystal structure of deoxycytidylate hydroxymethylase from bacteriophage T4, a component of the deoxyribonucleoside triphosphate-synthesizing complex. EMBO J. 18, 1104–1113.

Theobald, D.L., Mitton-Fry, R.M., and Wuttke, D.S. (2003). Nucleic acid recognition by OB-fold proteins. Annu. Rev. Biophys. Biomol. Struct. 32, 115.

Tian, H., Wu, Z., Chen, S., Ao, K., Huang, W., Yaghmaiean, H., Sun, T., Xu, F., Zhang, Y., and Wang, S. (2021). Activation of TIR signalling boosts pattern-triggered immunity. Nature 598, 500–503.

van Beljouw, S.P., Haagsma, A.C., Rodríguez-Molina, A., van den Berg, D.F., Vink, J.N., and Brouns, S.J. (2021). The gRAMP CRISPR-Cas effector is an RNA endonuclease complexed with a caspase-like peptidase. Science 373, 1349–1353.

Weijman, J.F., Kumar, A., Jamieson, S.A., King, C.M., Caradoc-Davies, T.T., Ledgerwood, E.C., Murphy, J.M., and Mace, P.D. (2017). Structural basis of autoregulatory scaffolding by apoptosis signal-regulating kinase 1. Proc. Natl. Acad. Sci. USA 114, E2096–E2105.

Wu, M., Sherwin, T., Brown, W.L., and Stockley, P.G. (2005). Delivery of antisense oligonucleotides to leukemia cells by RNA bacteriophage capsids. Nanomed. Nanotechnol. Biol. Med. 1, 67–76.

Yang, J., Roe, S.M., Cliff, M.J., Williams, M.A., Ladbury, J.E., Cohen, P.T., and Barford, D. (2005). Molecular basis for TPR domain-mediated regulation of protein phosphatase 5. EMBO J 24, 1–10.

Yu, G., Smith, D.K., Zhu, H., Guan, Y., and Lam, T.T.Y. (2017). ggtree: an R package for visualization and annotation of phylogenetic trees with their covariates and other associated data. Methods Ecol. Evol. 8, 28–36.

Zeytuni, N., and Zarivach, R. (2012). Structural and functional discussion of the tetra-trico-peptide repeat, a protein interaction module. Structure 20, 397–405.

Zhao, G., Wu, G., Zhang, Y., Liu, G., Han, T., Deng, Z., and He, X. (2014). Structure of the N-glycosidase MilB in complex with hydroxymethyl CMP reveals its Arg23 specifically recognizes the substrate and controls its entry. Nucleic Acids Res 42, 8115–8124.

Zhou, X., Liao, H., Chern, M., Yin, J., Chen, Y., Wang, J., Zhu, X., Chen, Z., Yuan, C., and Zhao, W. (2018). Loss of function of a rice TPR-domain RNA-binding protein confers broad-spectrum disease resistance. Proc. Natl. Acad. Sci. USA 115, 3174–3179.

